# In silico results of ϰ-Opioid receptor antagonists as ligands for the second bromodomain of the Pleckstrin Homology Domain Interacting Protein

**DOI:** 10.1101/432468

**Authors:** Lemmer R. P. El Assal

## Abstract

Pleckstrin Homology Domain Interacting Protein (PHIP) is a member of the BRWD1-3 Family (Bromodomain and WD repeat-containing proteins). PHIP (BRWD2, WDR11) contains a WD40 repeat (methyl-lysine binder) and 2 bromodomains (acetyl-lysine binder). It was discovered through interactions with the pleckstrin homology domain of Insulin Receptor Signalling (IRS) proteins and has been shown to mediate transcriptional responses in pancreatic islet cells and postnatal growth. An initial hit for the second bromodomain of PHIP (PHIP(2)) was discovered in 2012, with consecutive research yielding a candidate with a binding affinity of 68μM. PHIP(2) is an atypical category III bromodomain with a threonine (THR1396) where an asparagine residue would usually be. In the standard case, this pocket holds four water molecules, but in the case of PHIP(2), there is room for one extra water molecule - also known as “PHIP water”, able to mediate interaction between THR1396 and the typical water network at the back of the binding pocket. We present first ever results of two ϰ-Opioid receptor (KOR) antagonists with distinct pharmacophores having an estimated binding affinity in the nM to μM range, as well as higher binding affinities for every currently discovered PHIP(2) ligand towards KOR. Finally, we also demonstrate selectivity of LY-255582 and LY-2459989 towards PHIP(2) over other bromodomains.

## Introduction

Pre-administration of methadone was shown to enhance the apoptotic effect of doxorubicin in leukemia cells with a proposed mechanism involving lowered cAMP concentrations by inhibition of adenylyl cyclase, leading to increased caspase activity. Likewise, significant reduction in apoptosis was demonstrated following an increase in intracellular cAMP concentration[1, 2, 3] - it may however be possible that an increase in cAMP would simply mean an increase in CREBBP activity, a crucial transcriptional coactivator in the expression of P-Glycoprotein (P-GP) [4]. Furthermore, this hypothesis relies on the assumption that methadone is a P-GP inhibitor by itself, while it is only shown as a substrate[5, 6, 7], while other sources report significant reduction of methadone efflux with the specific by addition of P-GP inhibitors[8, 9, 10]. P-GP expression is directly regulated by attachment of a variety of transcriptional factors to the promoter region of the P-GP gene; examples thereof are: p53[11], YB-1[12] and NF-*κ*B[13].

Bromodomains are acetyl-lysine (KAc) reader regions involved in the modulation of gene expression.[14]They have now for a while been attractive targets for the treatment of diseases such as inflammation and cancer, bringing about the development of a range of chemical probes for the investigation of bromodomain (Brd) biology.[15] In 2010, JQ-1 and I-BET were reported as potent inhibitors of the bromo- and extra-terminal domain (BET) bromodomain,[16, 17]and later academic and industrial research has been targeting the BET bromodomains.[18, 19, 20, 21]

Pleckstrin Homology Domain Interacting Protein (PHIP) is a member of the BRWD1-3 Family (Bromodomain and WD repeat-containing proteins). PHIP (BRWD2, WDR11) contains a WD40 repeat (methyl-lysine binder) and 2 bromodomains (acetyl-lysine binder). It was discovered through interactions with the pleckstrin homology domain of Insulin Receptor Signalling (IRS) proteins and has been shown to mediate transcriptional responses in pancreatic islet cells and postnatal growth.

In addition to its role in IGF-signalling, it has been identified as the gene most highly overexpressed in metastatic melanomas, compared with primary tumors[22], activation of PHIP promotes melanoma metastasis, and can be used as a biomarker to classify a subset of primary melanomas[23], and that elevated PHIP copy number was associated with significantly reduced distant metastasis-free survival and disease-specific survival by Kaplan-Meier analyses and that PHIP plays an important role as a molecular marker of melanoma ulceration, metastasis and survival[24]. Furthermore, it was identified as an increased risk in breast cancer[25] and as a tumour suppressor in Group 3 medulobalstoma [26].

An initial hit for the second bromodomain of PHIP (PHIP(2)) was discovered in 2012 (PDB 3MB3[14]), with consecutive research yielding a candidate with a binding affinity of 68μM[27, 28]. PHIP(2) is an atypical category III bromodomain[14] with a threonine (THR1396) where an asparagine residue would usually be. In the standard case, this pocket holds four water molecules, but in the case of PHIP(2), there is room for one extra water molecule - also known as “PHIP water”, able to mediate interaction between THR1396 and the typical water network at the back of the binding pocket.

Seeing how insulin can induce expression of P-GP[13], and PHIP is involved in insulin signaling, we decided to dock methadone against PHIP(2) to evaluate whether or not methadone in fact works by inhibiting interaction with KAc residues of histones by attaching to bromodomains - more specifically - PHIP(2). Upon estimating binding affinity in the micromolar region for methadone against PDB 5ENB[27] using HYDE [29, 30], a random set of opioids was chosen for screening. With more data, the search has shown a variety of ϰ-Opioid receptor (KOR) antagonists having good estimated binding affinities against PHIP(2). Finally, two candidates with distinct pharmacophores were selected to probe against other bromodomains to show selectivity towards PHIP(2). Additionally, it was shown that every single molecule with high reported affinity towards PHIP(2) is estimated to have even higher affinity towards KOR.

### Results and discussion

#### Initial search

The initial search was not very successful, yielding only two hits with nanomolar affinity. Even though MCL-117 shows no H-bonds towards the target, it was decided that the selectivity of both compounds towards μ and ϰ in comparison to δ warranted a restriction from including δ-Opioid receptor ligands from further screening.

**Table 1:**
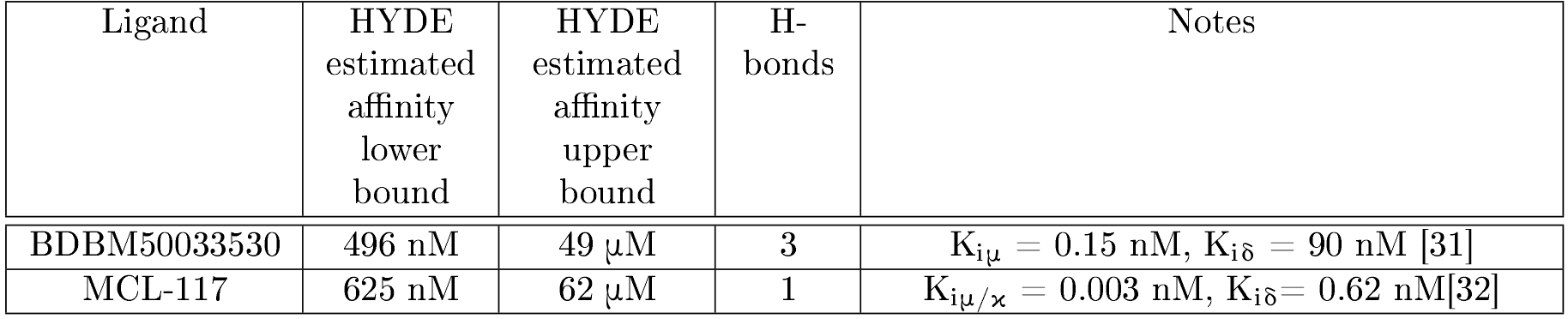
Results for an initial search

| Ligand | HYDE<br>estimated<br>affinity<br>lower<br>bound | HYDE<br>estimated<br>affinity<br>upper<br>bound | H-bonds | Notes |
| --- | --- | --- | --- | --- |
| BDBM50033530 | 496 nM | 49 μM | 3 | $K_{i\mu} = 0.15$ nM, $K_{i\delta} = 90$ nM [31] |
| MCL-117 | 625 nM | 62 μM | 1 | $K_{i\mu/\kappa} = 0.003$ nM, $K_{i\delta} = 0.62$ nM[32] |

#### ϰ-subtype agonists

Even though the results are not that astonishing (since the estimated binding affinity range is not comparable with BDBM50033530), Cyclorphan has an estimated binding affinity that about ten-fold that of the experimental value of FMOOA463, although with only one recognized H-bond. A potentially additional residue could form an H-bond with the nitrogen of the piperidine substructure: Tyr1350 (fig 1). This is unfortunately not confirmed because it is only recognized as hydrophobic interaction. In the case of Nalfurafine (fig 2), even though there is a predicted H-bond with THR1396, the estimated binding affinity range is in the mM region and the uncommon torsions make it unattractive for further investigation.

**Figure 1:**
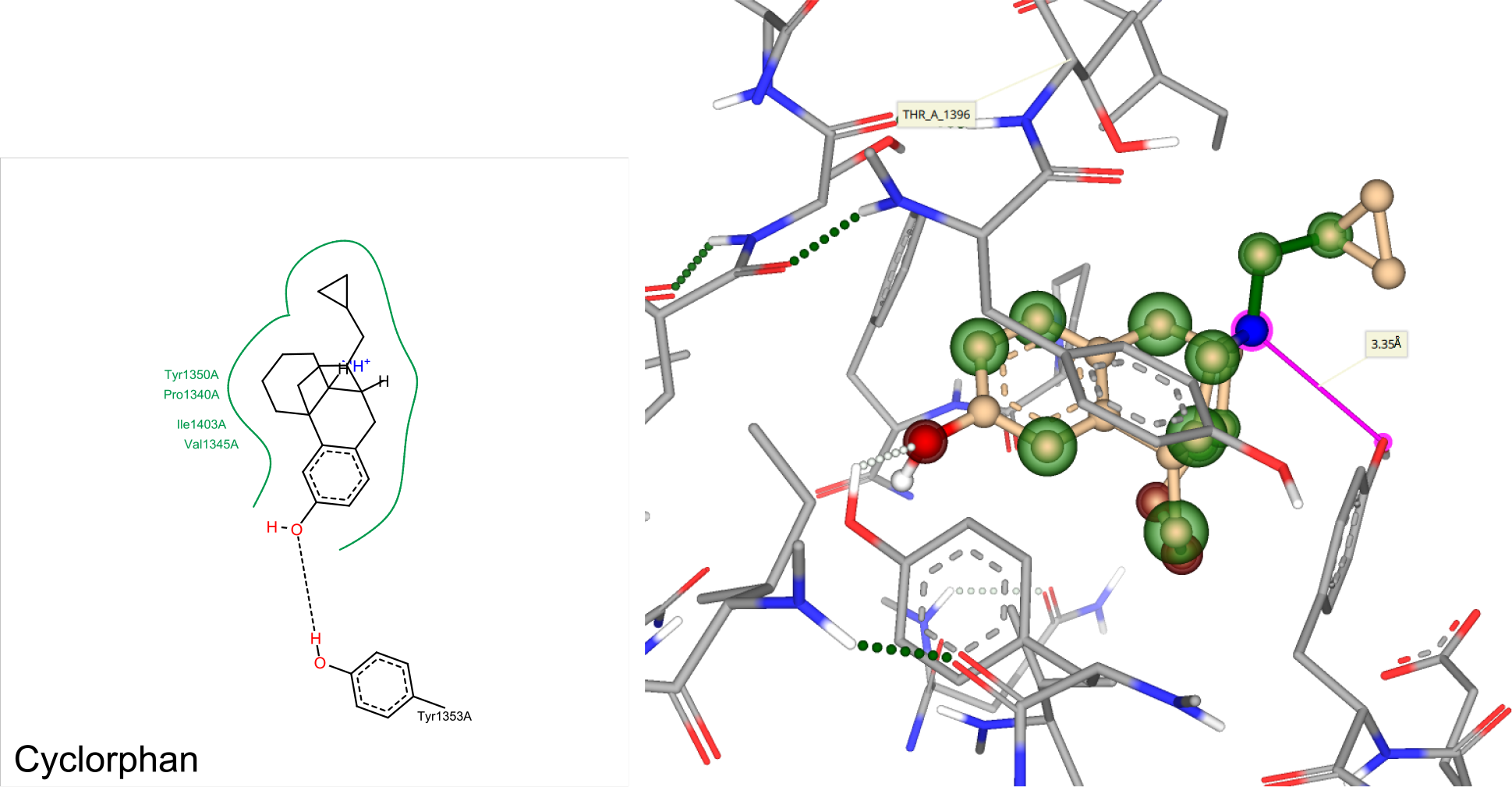
Three-dimensional docking view of Cyclorphan in SeeSAR

**Figure 2:**
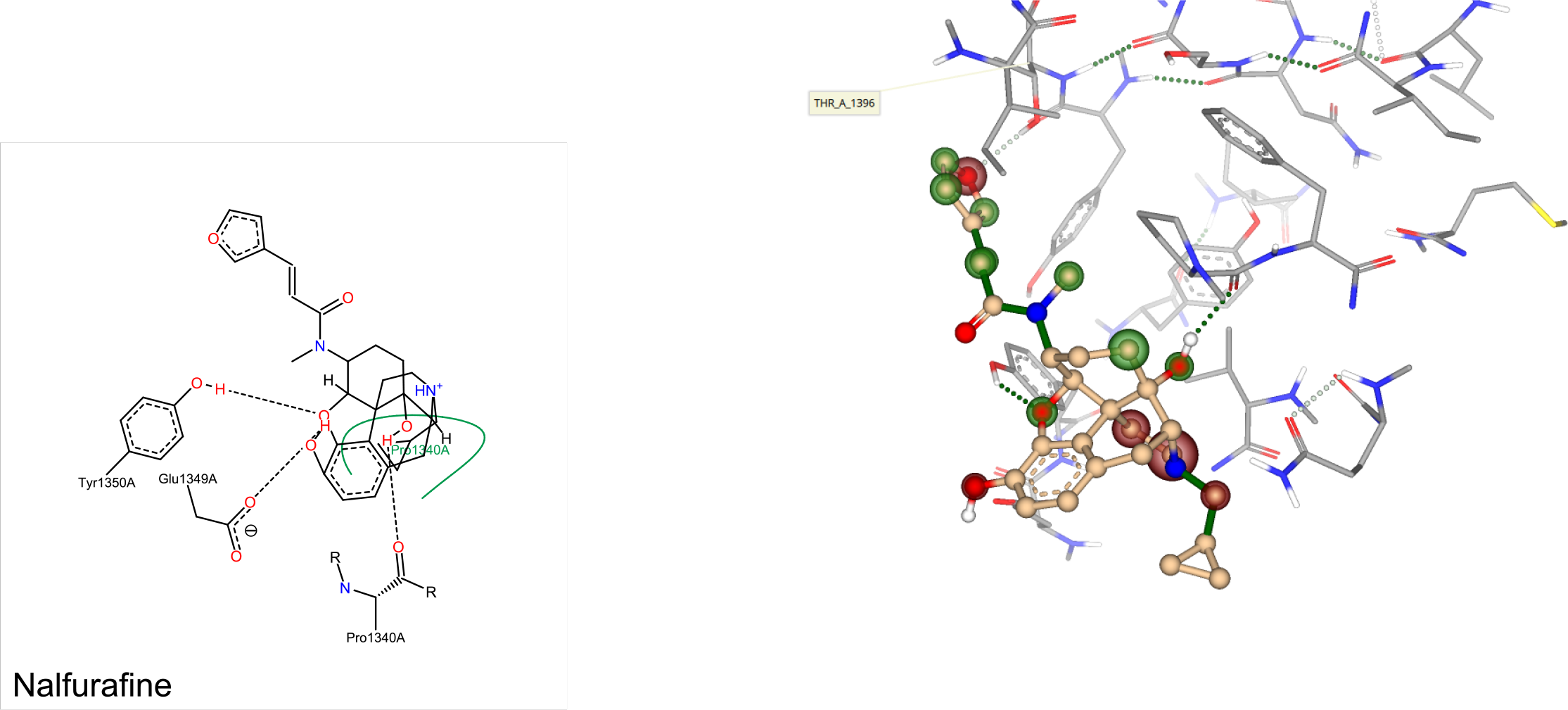
Three-dimensional docking view of Nalfurafine in SeeSAR

**Table 2:**
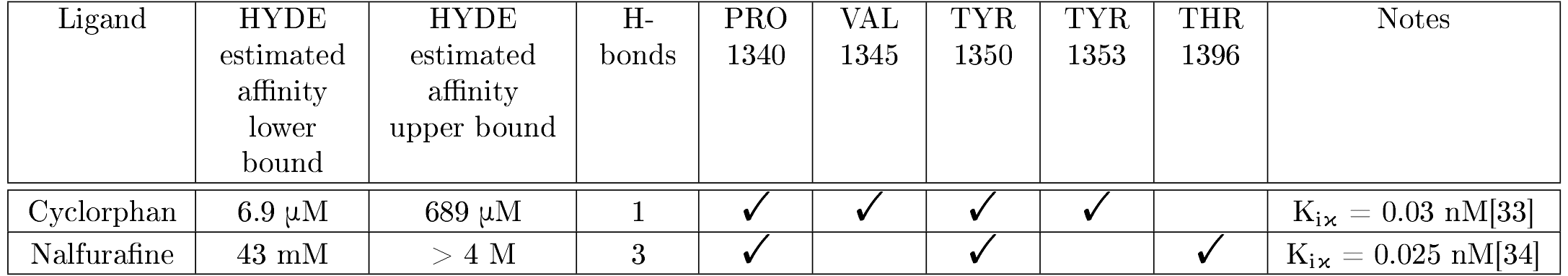
Results for docking of KOR agonists

#### ϰ-subtype antagonists

Since no ligand bearing green torsions with better binding affinity could be found over BDBM50033530, no definitive conclusion could be made and thus, the search continued on the ϰ-subtype antagonists instead.

**Table 3:**
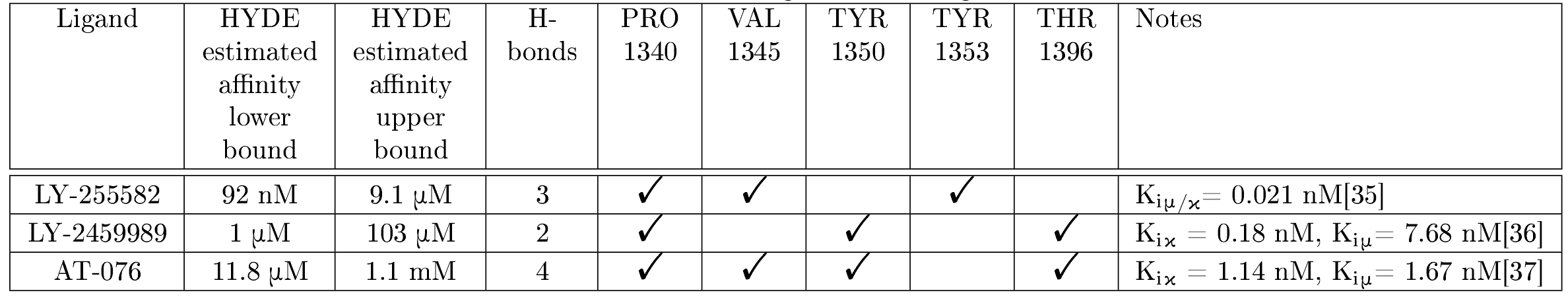
Results for docking of KOR antagonists

A ligand with much higher estimated binding affinity was discovered which, like BDBM50033530, bears 3 H-bonds towards the target (fig 3). Seeing how the target is to stop PHIP(2) from interacting with acetyl-lysine residues, it would make sense that, should there be any link whatsoever between PHIP(2) and KOR, antagonists would have better binding affinity, explaining the better binding affinity estimations.

**Figure 3:**
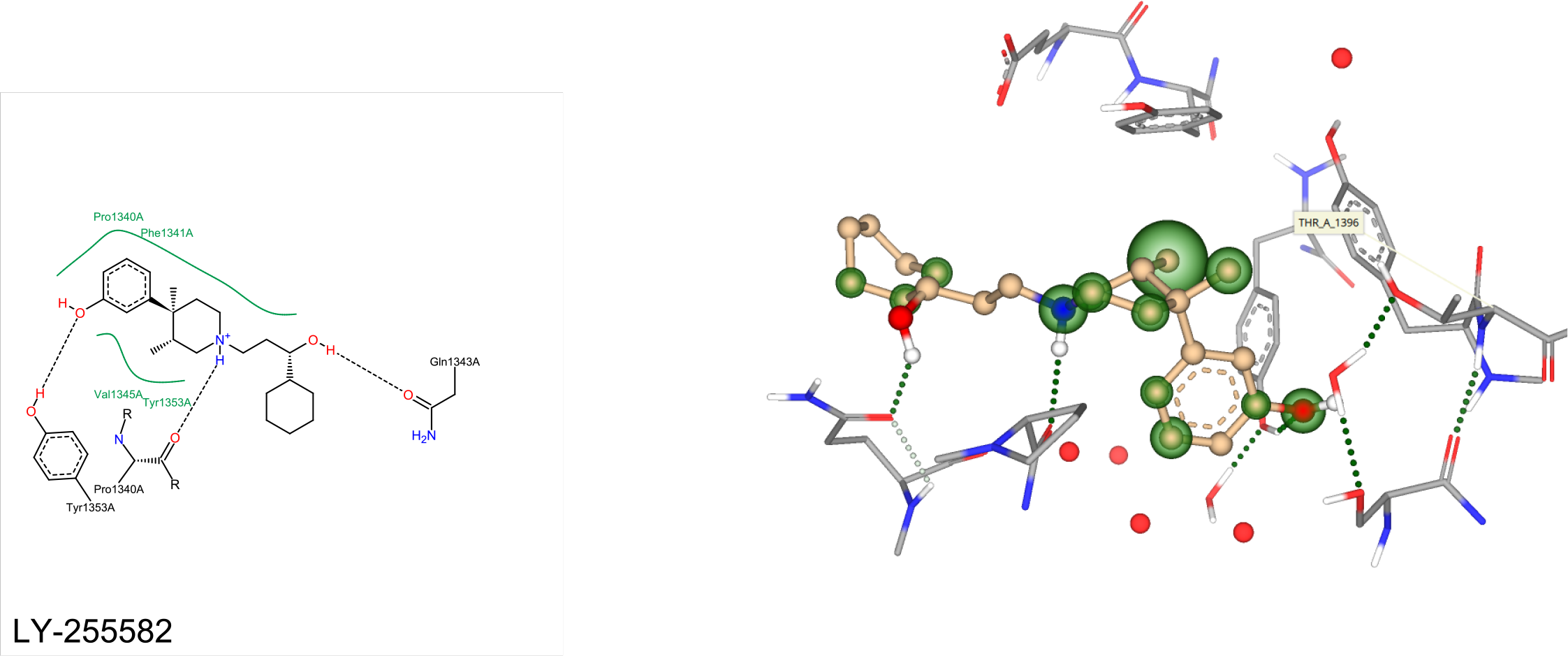
Three-dimensional docking view of LY-255582 in SeeSAR

In the case of LY-2459989 (fig 4) and AT-076 (fig 5), there is an H-bond network with THR1396.

**Figure 4:**
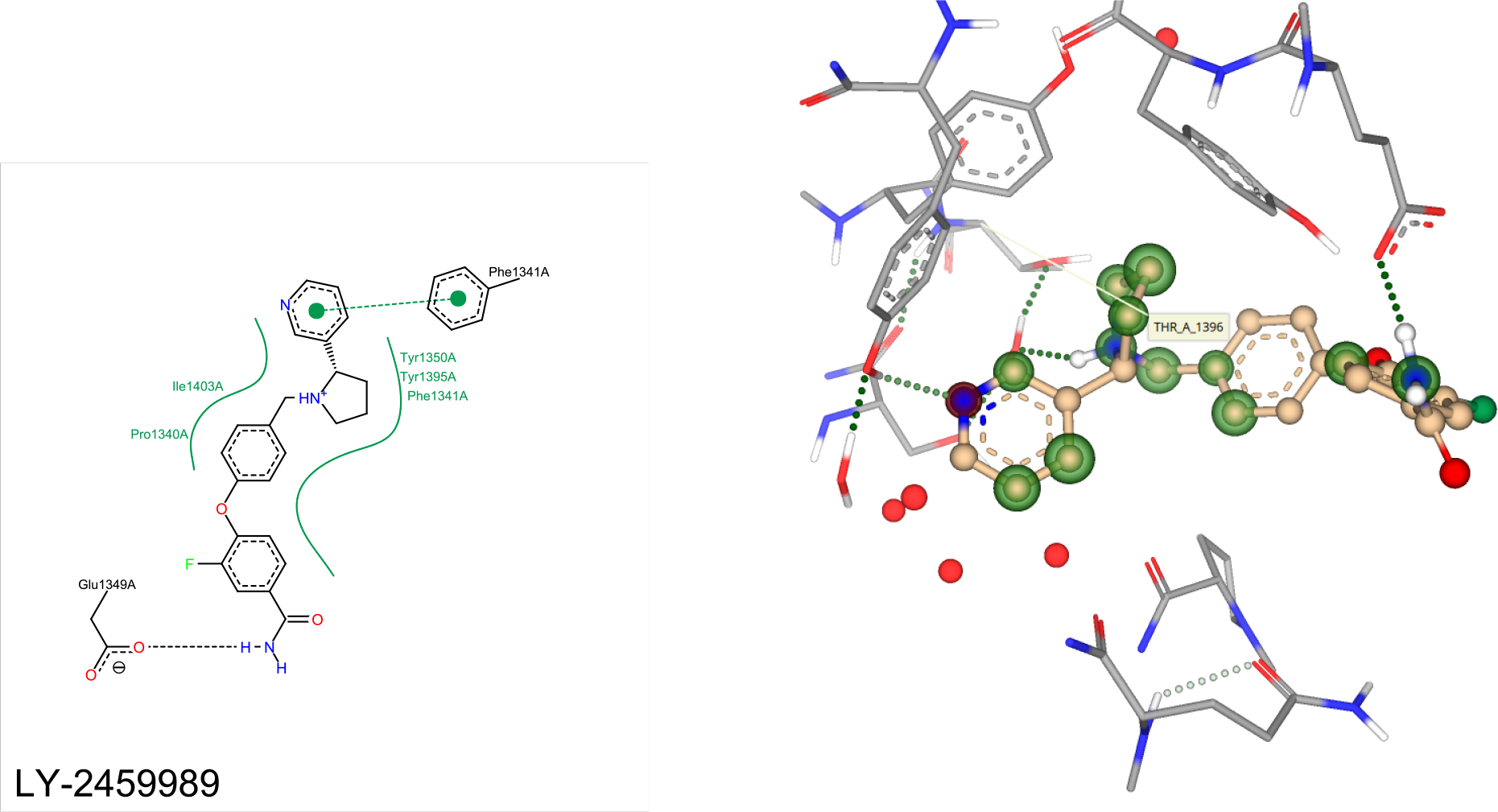
Three-dimensional docking view of LY-2459989 in SeeSAR

**Figure 5:**
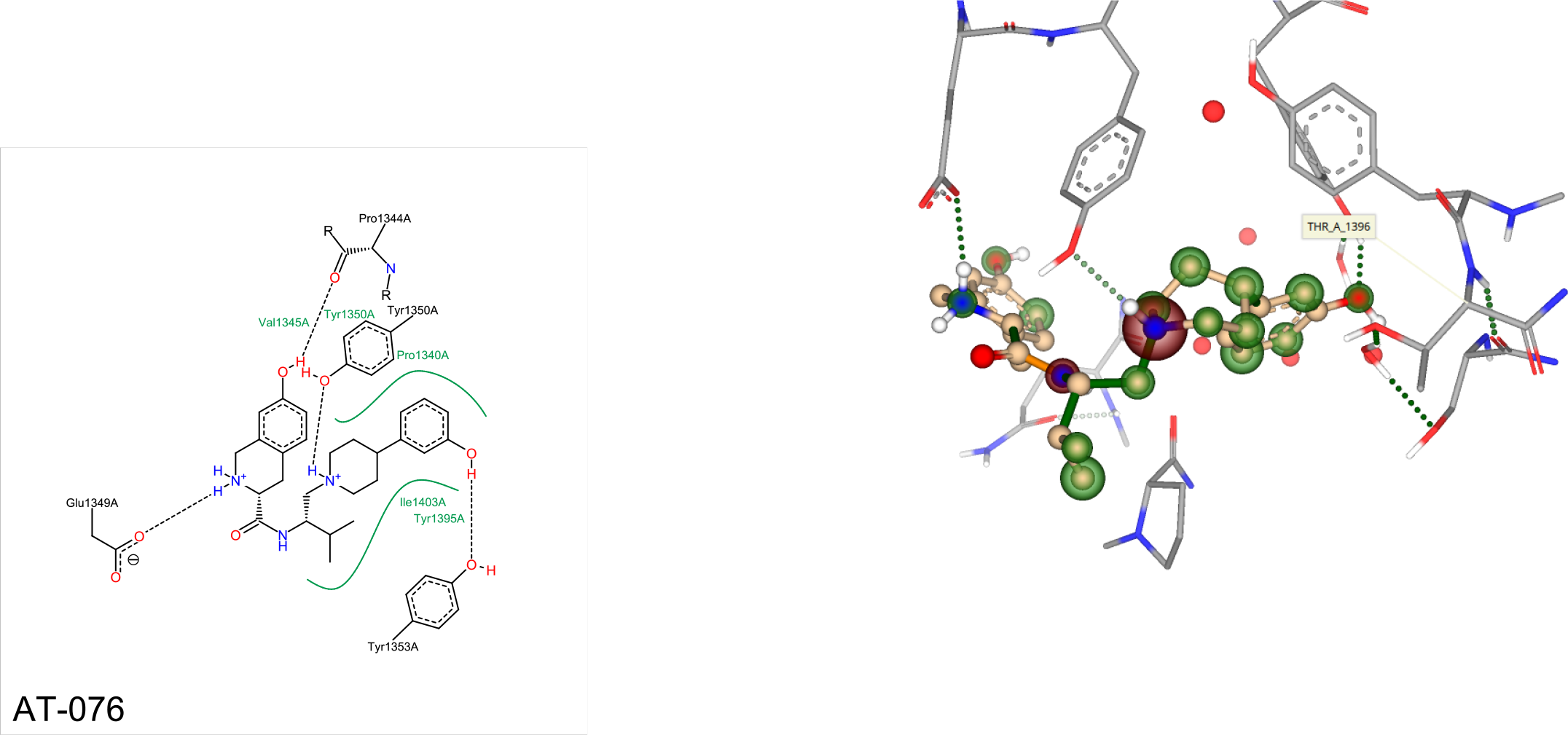
Three-dimensional docking view of AT-076 in SeeSAR

#### Docking of LY-2459989 and LY-255582 against EP300

The estimated affinities are similar to PHIP(2) for both ligands. With three H-bonds being exclusively one-sided and no π-interaction to hold it in place, the result is unreliable for LY-2459989. LY-255582, on the other hand, shows a solid support from both sides using H-bonds. This is unsurprising, as EP300 and PHIP(2) share the same category[14].

**Table 4:** Docking of LY-2459989 and LY-255582 against EP300

| Ligand | HYDE<br>estimated<br>affinity lower<br>bound | HYDE<br>estimated<br>affinity upper<br>bound | H-bonds |
| --- | --- | --- | --- |
| LY-2459989 | 82 nM | 8.2 μM | 3 |
| LY-255582 | 1.3 μM | 125 μM | 2 |

**Figure 6:**
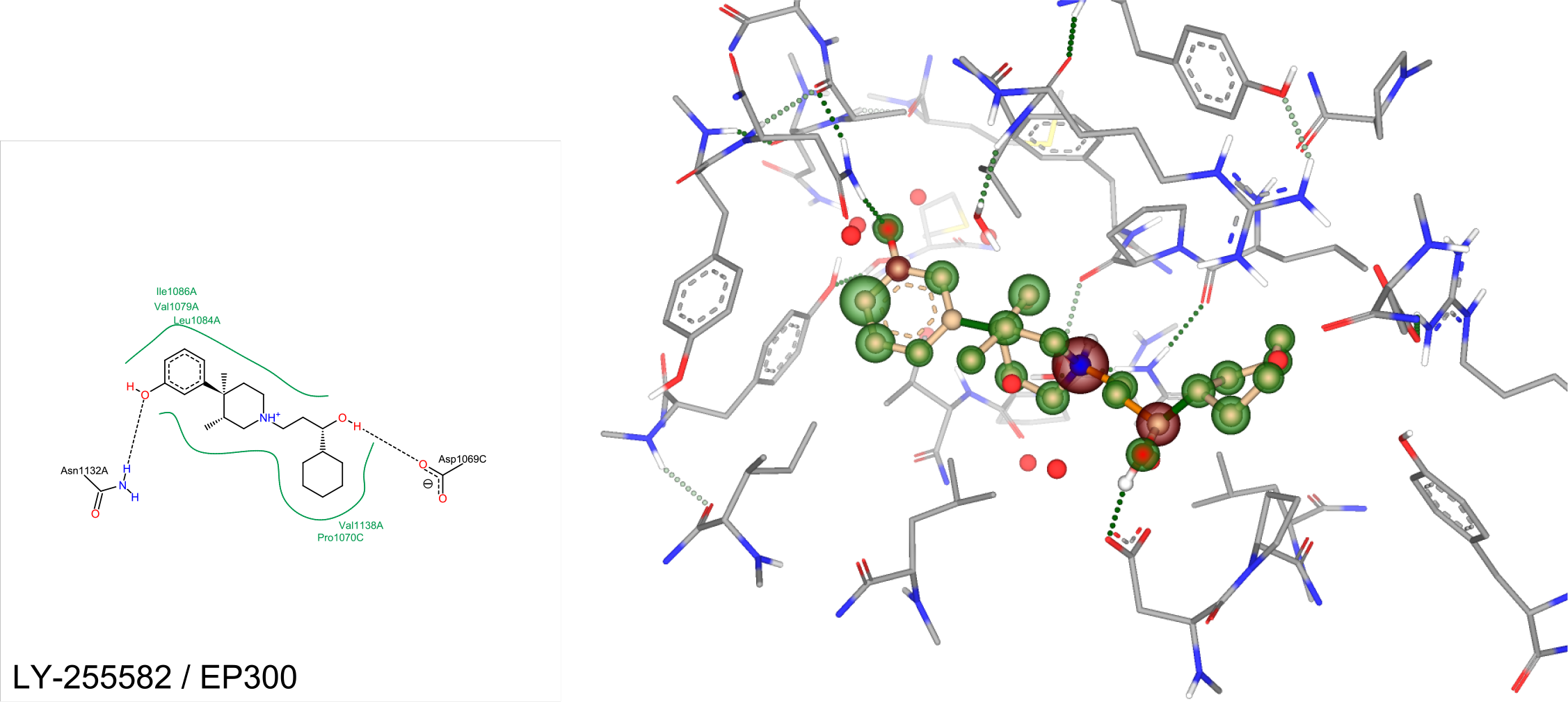
Three-dimensional docking view of LY-255582 against EP300 in SeeSAR

**Figure 7:**
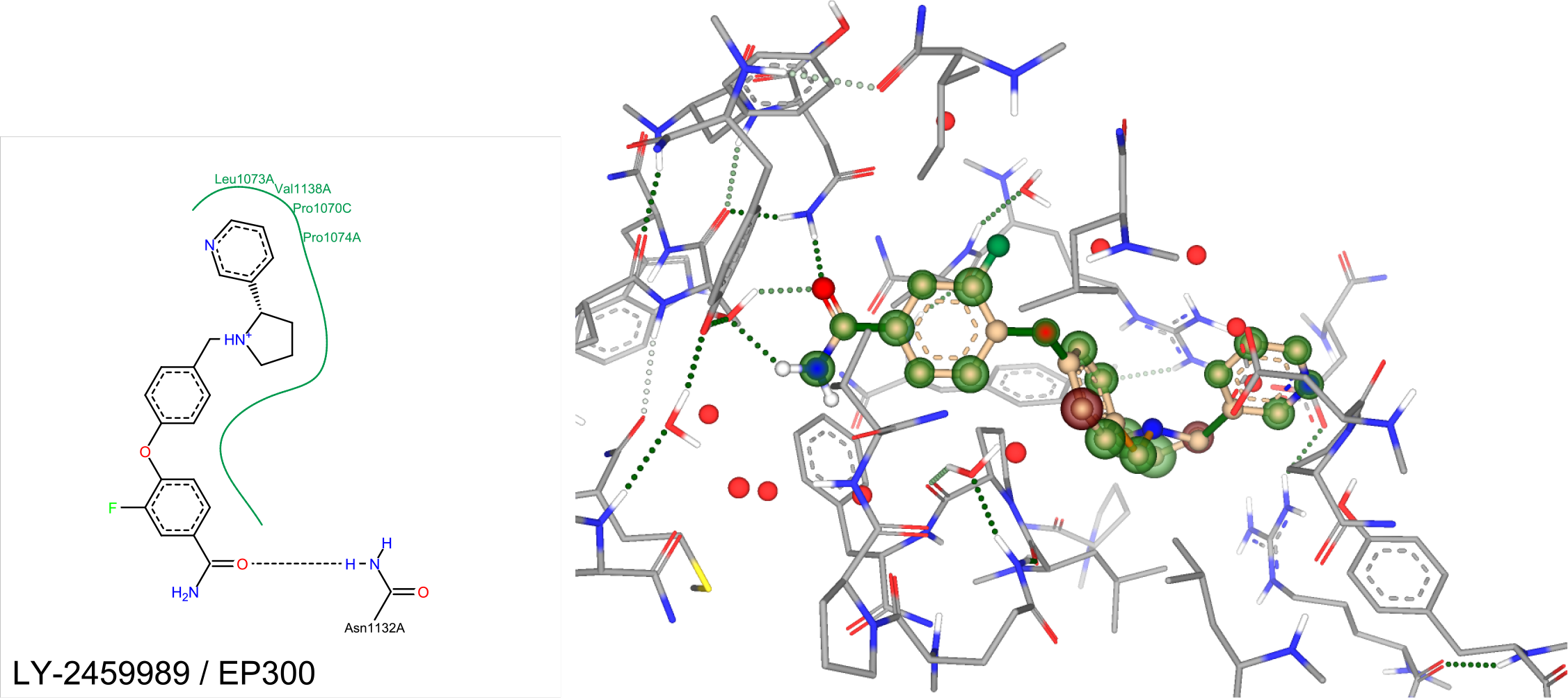
Three-dimensional docking view of LY-2459989 against EP300 in SeeSAR

#### Docking of LY-2459989 and LY-255582 against BRD9

The estimated affinity for both ligands is about half as good as for PHIP(2). LY-255582 is supported from both sides (notably by π-interaction from Tyr106A), although only using 2 H-bonds (rather than 3 in the case of PHIP(2)). In the case of LY-2459989, there is only one H-bond, making it very unbalanced, even though it there is π-interaction from Tyr106A. This is good, because BRD9 is not in the same category as PHIP(2)[14].

**Table 5:** Docking of LY-2459989 and LY-255582 against BRD9

| Ligand | HYDE<br>estimated<br>affinity lower<br>bound / nM | HYDE<br>estimated<br>affinity upper<br>bound / nM | Torsion colour | H-bonds |
| --- | --- | --- | --- | --- |
| LY-255582 | 199 | 19 778 | Green | 2 |
| LY-2459989 | 1 981 | 196 853 | Green | 1 |

**Figure 8:**
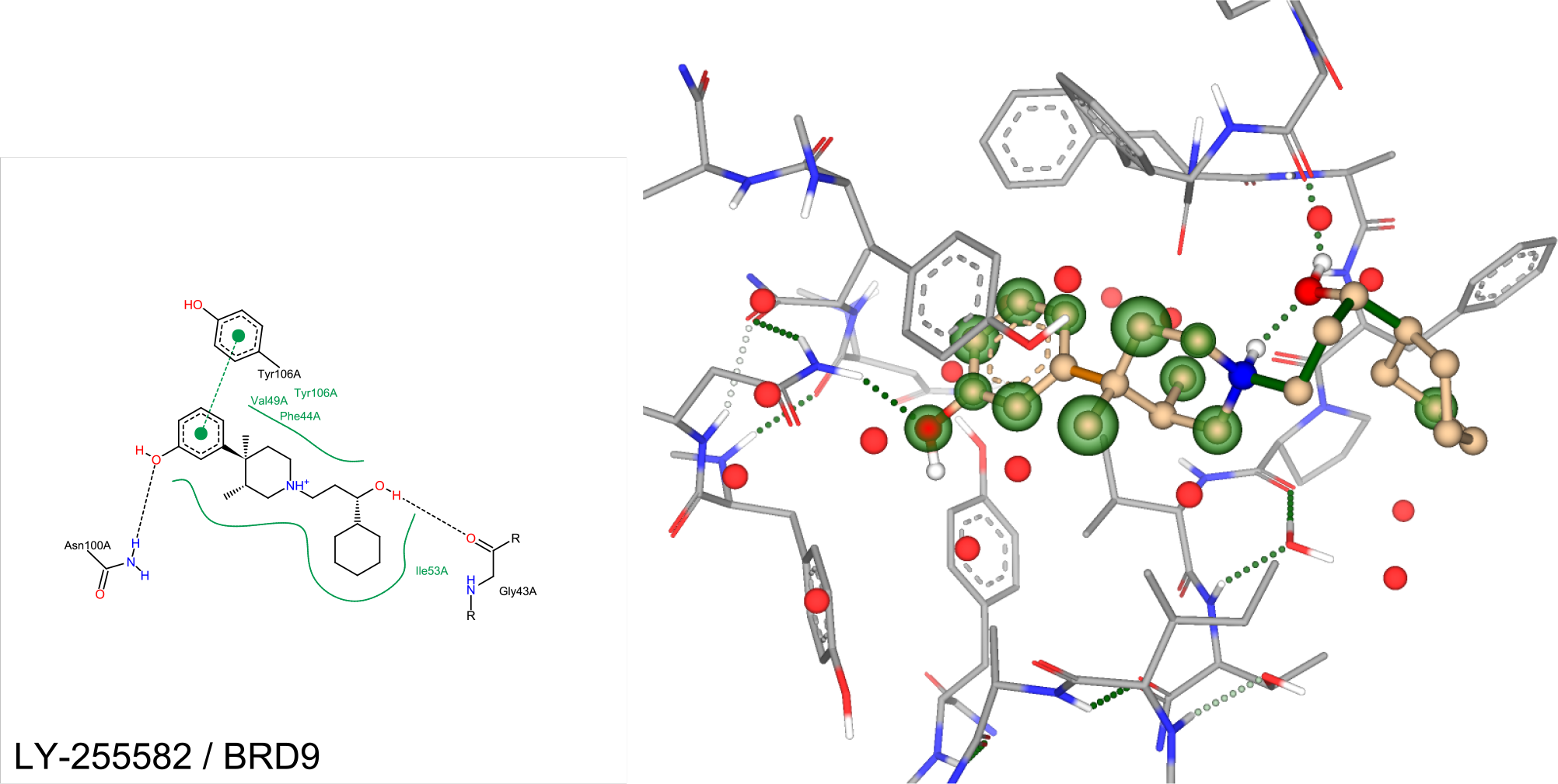
Three-dimensional docking view of LY-255582 against BRD9 in SeeSAR

**Figure 9:**
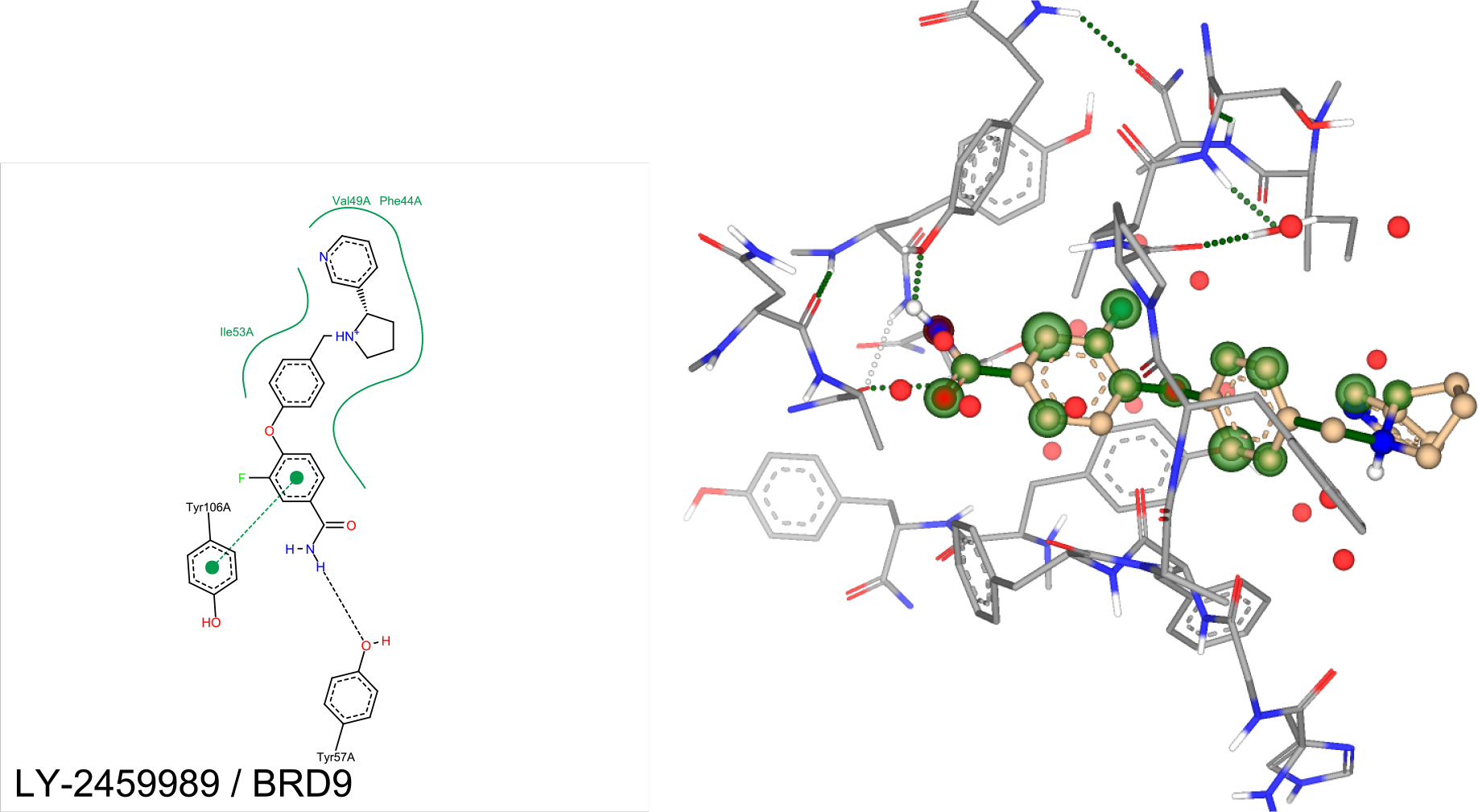
Three-dimensional docking view of LY-2459989 against BRD9 in SeeSAR

#### Docking of LY-2459989 and LY-255582 against BRDI

The lower bound for the estimated affinity of LY-255582 is 290x weaker, and, LY-2459989 has a lower bound that’s 440x weaker in comparison with the estimated lower bound for PHIP(2)[14]. Both candidates are well supported by 3 H-bonds and LY-2459989 has additional π-interaction from TYR109A. This is very good, because it shows selectivity towards PHIP(2).

**Table 6:** Docking of LY-2459989 and LY-255582 against BRD1

| Ligand | HYDE<br>estimated<br>affinity lower<br>bound | HYDE<br>estimated<br>affinity upper<br>bound | H-bonds |
| --- | --- | --- | --- |
| LY-255582 | 26.8 μM | 2.7 mM | 3 |
| LY-2459989 | 459 μM | 45.6 mM | 3 |

**Figure 10:**
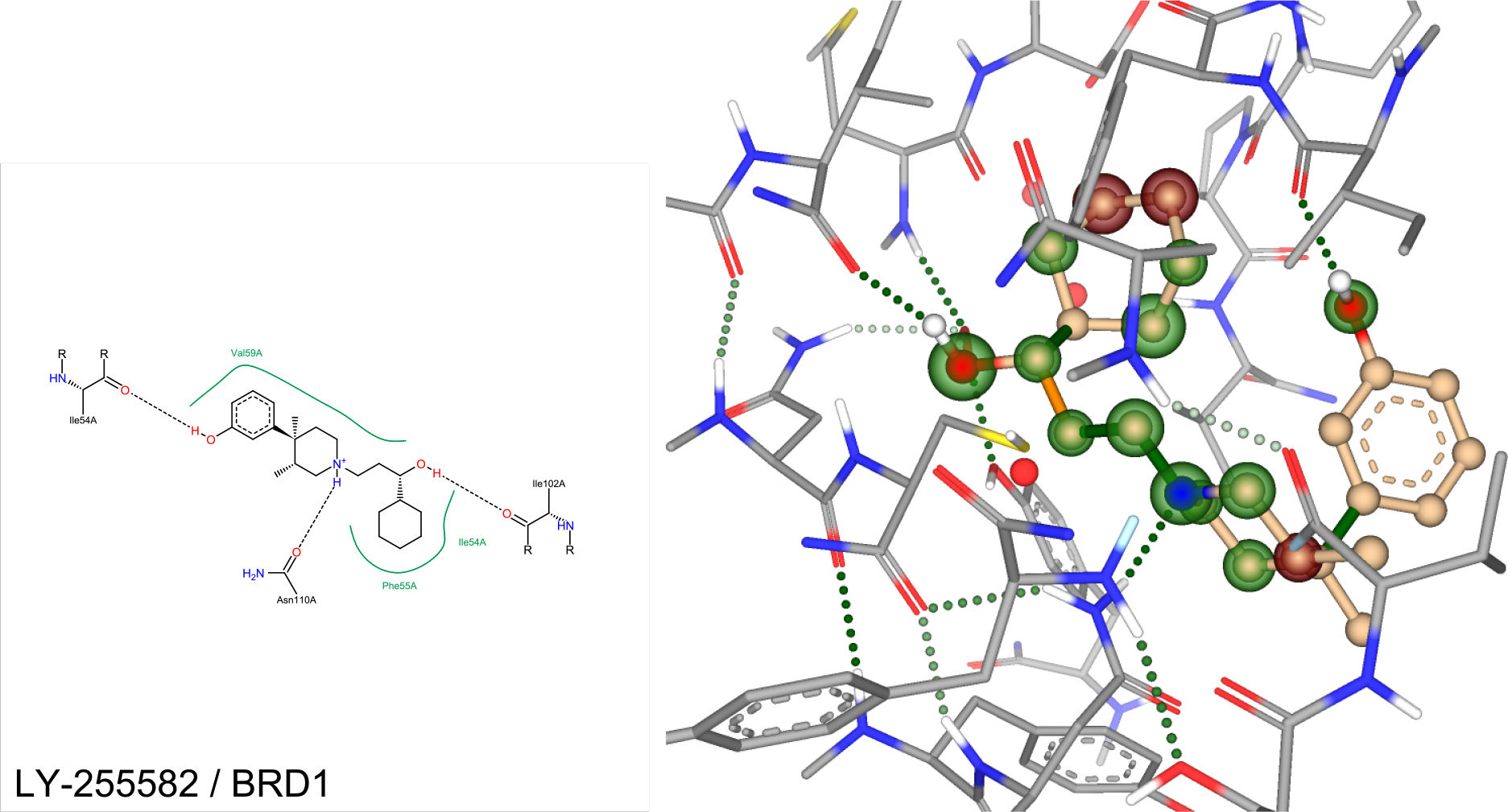
Three-dimensional docking view of LY-255582 against BRD1 in SeeSAR

**Figure 11:**
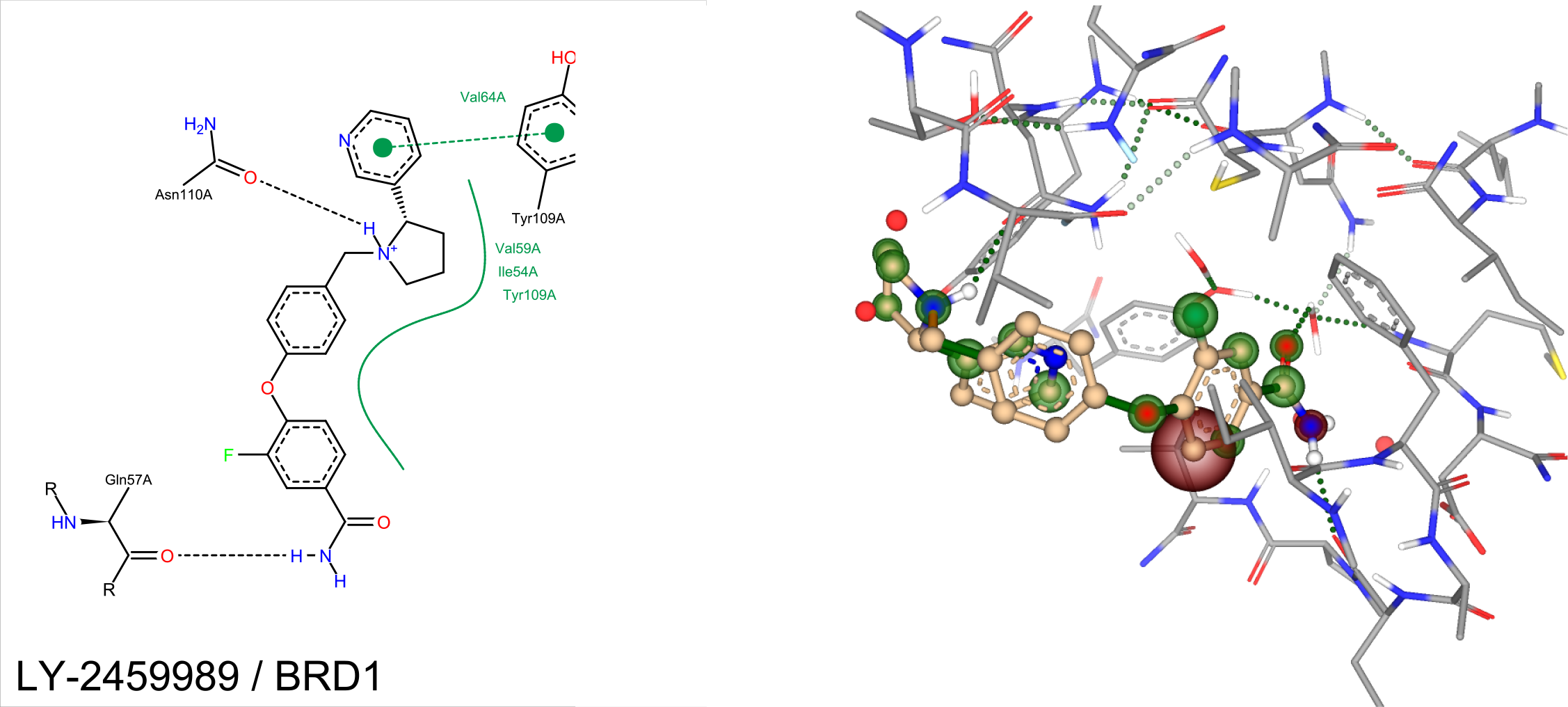
Three-dimensional docking view of LY-2459989 against BRD1 in SeeSAR

##### 0.1 Docking of LY-2459989 and LY-255582 against BRD7

The estimated affinities for LY-255582 is about 7 times weaker in comparison to PHIP(2) using 3 H-bonds and 2 π-interaction from Phe155A and Tyr217A. The estimated affinity for LY-2459989 is 1.6x higher towards BRD7 in comparison to PHIP(2), although using 2 balanced H-bonds and 1 π-interaction from TYR217A. Again, just like BRD9, these results are expected as PHIP(2) and BRD7 are not in the same category of bromodomains[14].

**Table 7:** Docking of LY-2459989 and LY-255582 against BRD7

| Ligand | HYDE<br>estimated<br>affinity lower<br>bound | HYDE<br>estimated<br>affinity upper<br>bound | H-bonds |
| --- | --- | --- | --- |
| LY-255582 | 648 nM | 64.4 μM | 2 |
| LY-2459989 | 654 nM | 65 μM | 3 |

**Figure 12:**
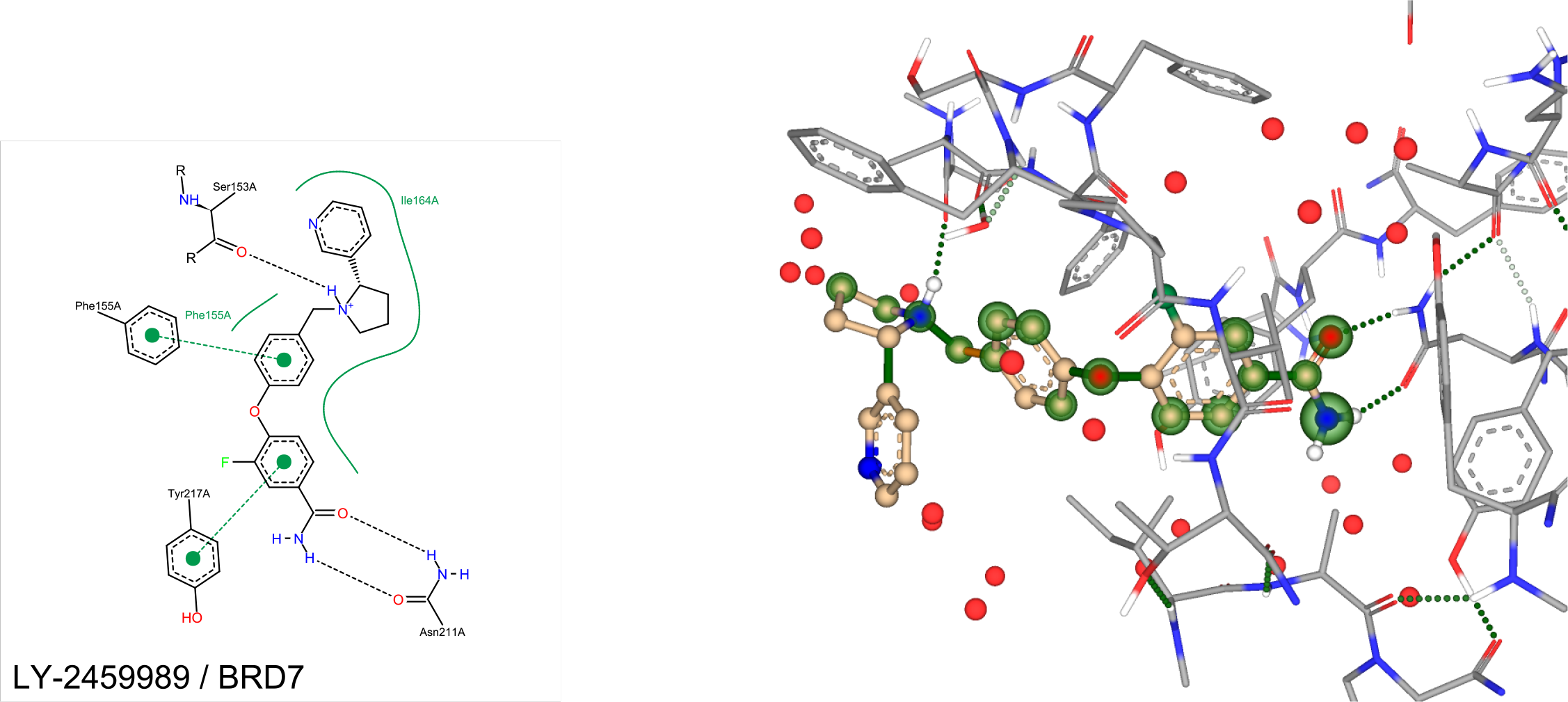
Three-dimensional docking view of LY-255582 against BRD7 in SeeSAR

**Figure 13:**
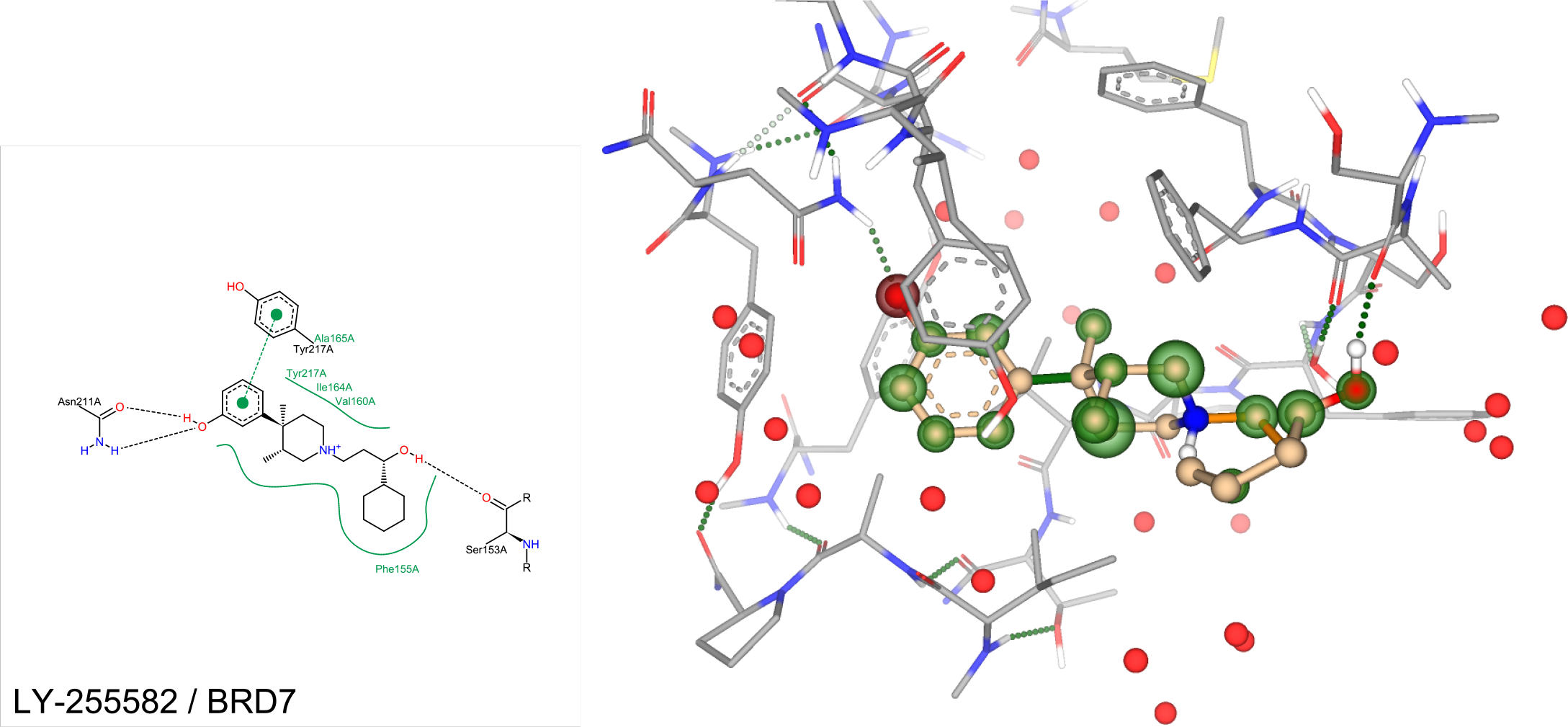
Three-dimensional docking view of LY-2459989 against BRD7 in SeeSAR

##### 0.2 Docking of LY-2459989 and LY-255582 against BRPFI

For LY-2459989, both H-bonds and π-interaction (with Phe714B) are one-sided, providing no good scaffolding support, similar to the results towards EP300. This is not the case for LY-255582: 3 H-bonds, as well as a π-interaction with Phe714B. The binding affinities are more selective towards PHIP(2) than BRPF1. This selectivity is wanted, because BRPF1 is category IV and PHIP(2) is category III.[14]

**Table 8:** Docking of LY-2459989 and LY-255582 against BRPF1

| Ligand | HYDE<br>estimated<br>affinity lower<br>bound | HYDE<br>estimated<br>affinity upper<br>bound | H-bonds |
| --- | --- | --- | --- |
| LY-255582 | 2.6 μM | 262 μM | 3 |
| LY-2459989 | 10.4 μM | 1 mM | 2 |

**Figure 14:**
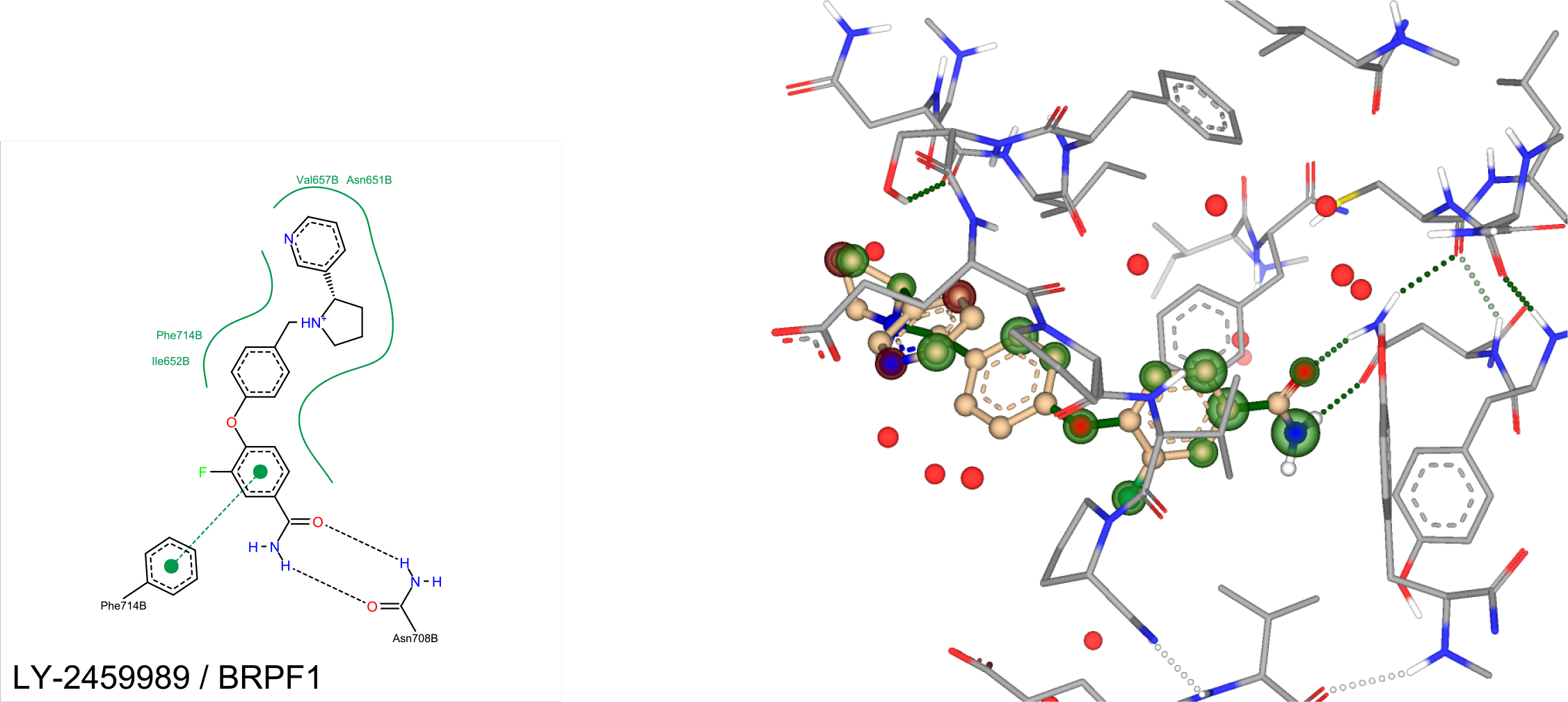
Three-dimensional docking view of LY-255582 against BRPF1 in SeeSAR

**Figure 15:**
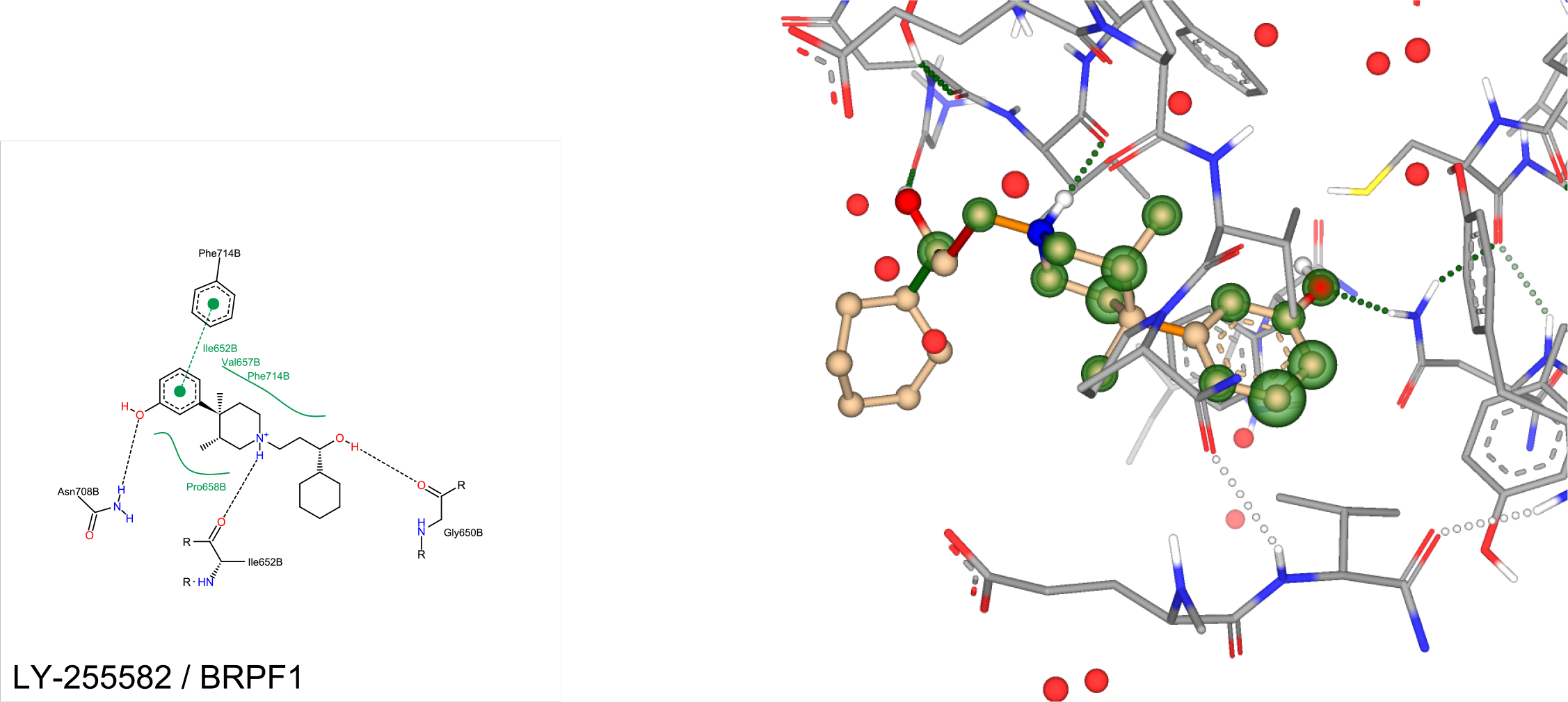
Three-dimensional docking view of LY-2459989 against BRPF1 in SeeSAR

##### 0.3 Docking of LY-2459989 and LY-255582 against CREBBP

The binding affinities for both are not that great in comparison to PHIP(2), even though they are in the same category of bromodomains[14].

**Table 9:**
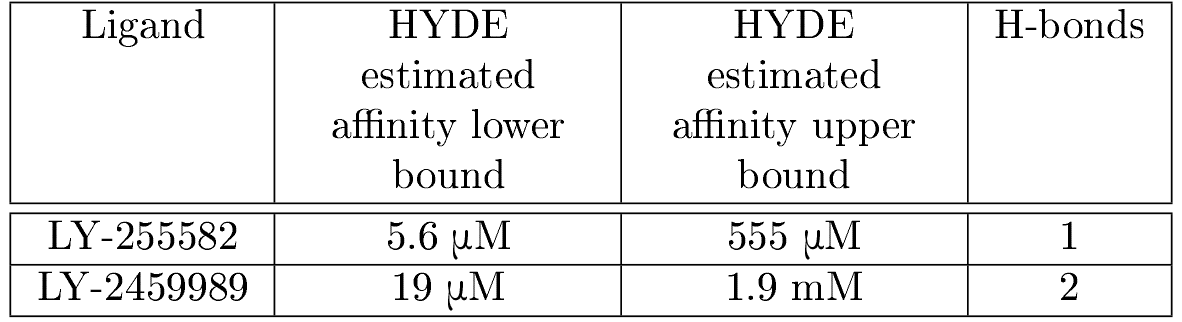
Docking of LY-2459989 and LY-255582 against CREBBP

| Ligand | HYDE<br>estimated<br>affinity lower<br>bound | HYDE<br>estimated<br>affinity upper<br>bound | H-bonds |
| --- | --- | --- | --- |
| LY-255582 | 5.6 μM | 555 μM | 1 |
| LY-2459989 | 19 μM | 1.9 mM | 2 |

**Figure 16:**
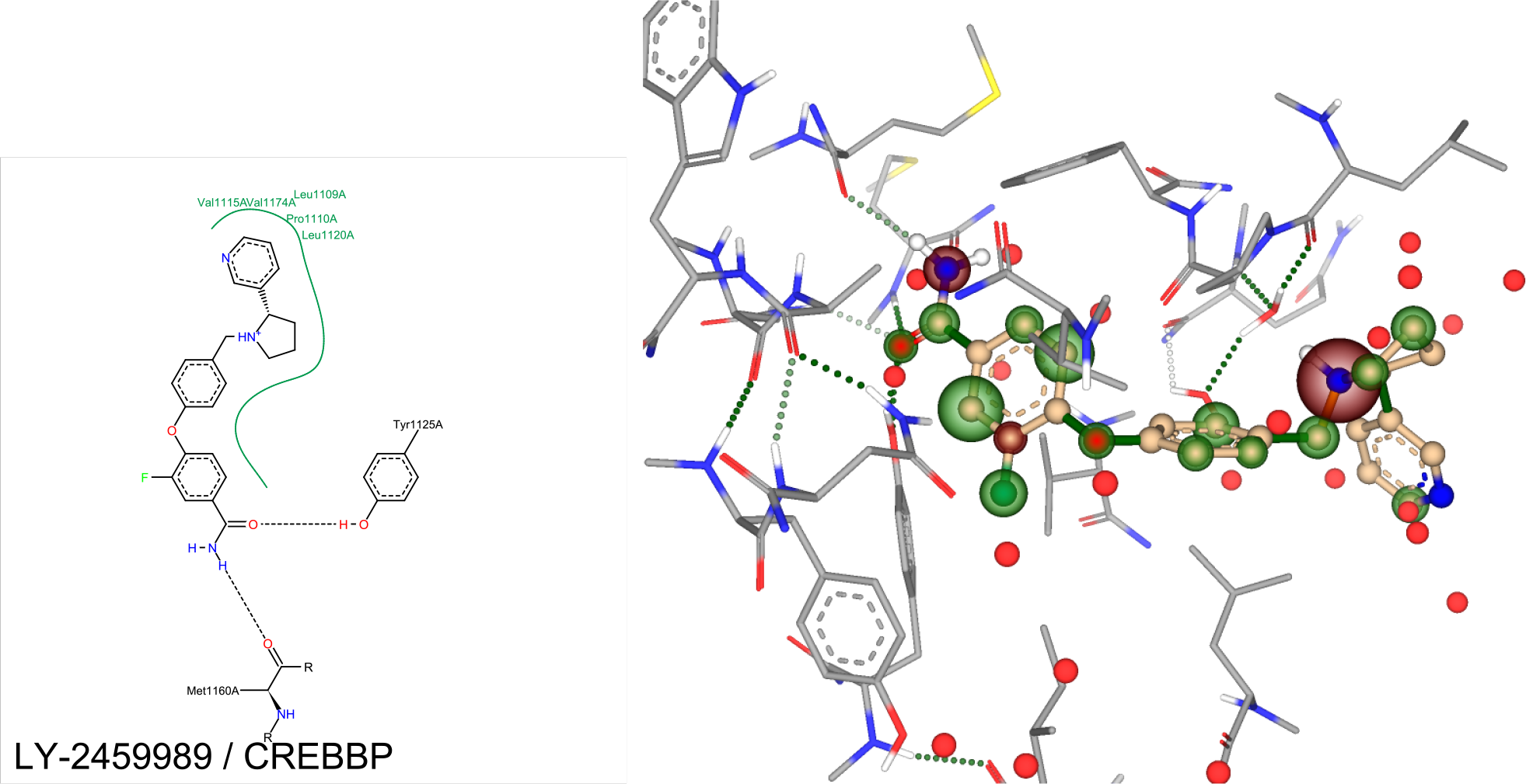
Three-dimensional docking view of LY-255582 against CREBBP in SeeSAR

**Figure 17:**
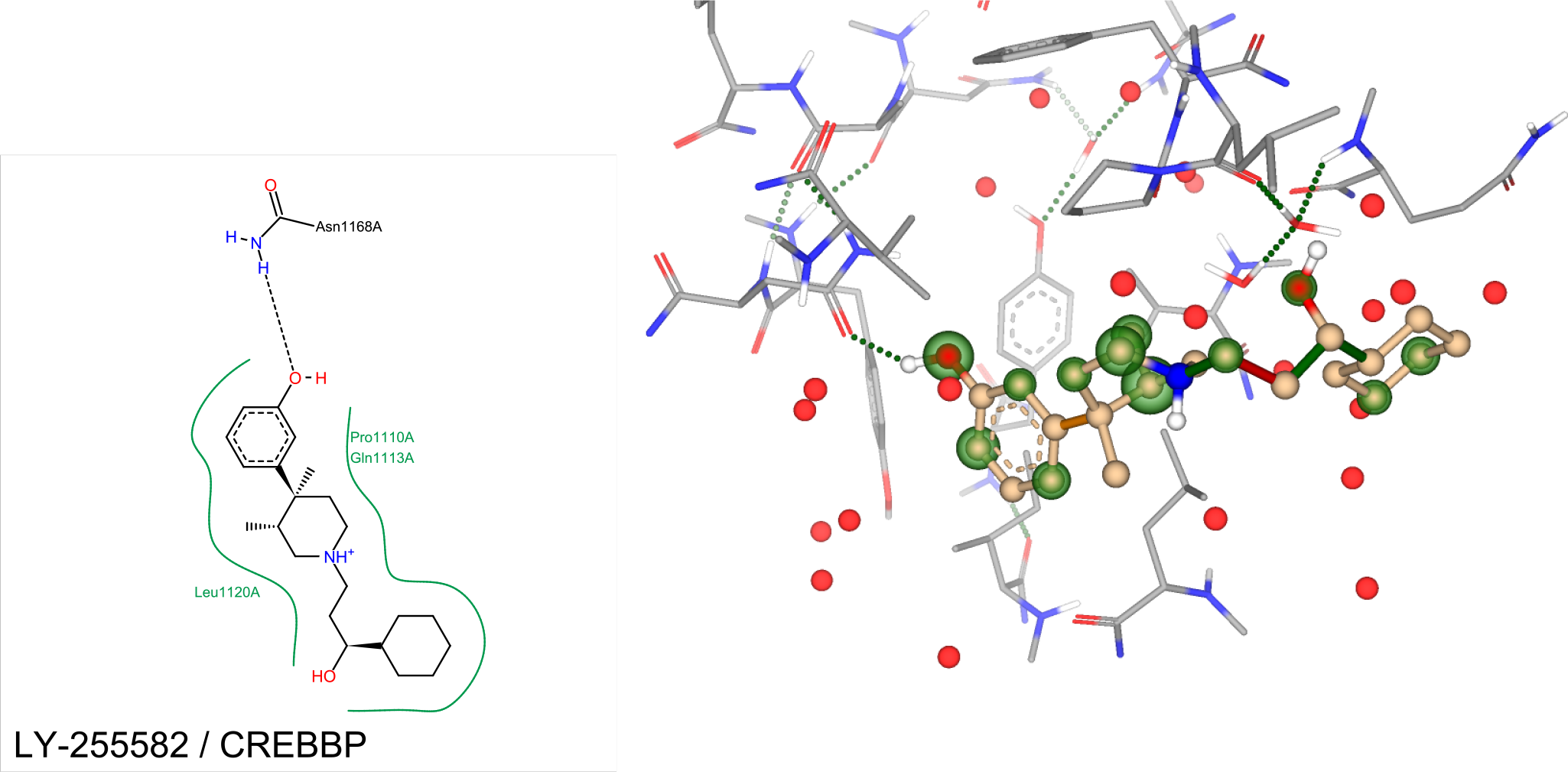
Three-dimensional docking view of LY-2459989 against CREBBP in SeeSAR

#### Docking of LY-2459989 and LY-255582 against BAZ2A

In both cases, 2 H-bonds hold the molecules in place and the estimated binding affinities are over 250x lower than for PHIP(2). This selectivity is advantageous, seeing how PHIP(2) is category III, whereas BAZ2A is category V[14].

**Table 10:** Docking of LY-2459989 and LY-255582 against BAZ2A

| Ligand | HYDE<br>estimated<br>affinity lower<br>bound | HYDE<br>estimated<br>affinity upper<br>bound | H-bonds |
| --- | --- | --- | --- |
| LY-255582 | 24 μM | 2.4 mM | 3 |
| LY-2459989 | 289 μM | 28 mM | 3 |

**Figure 18:**
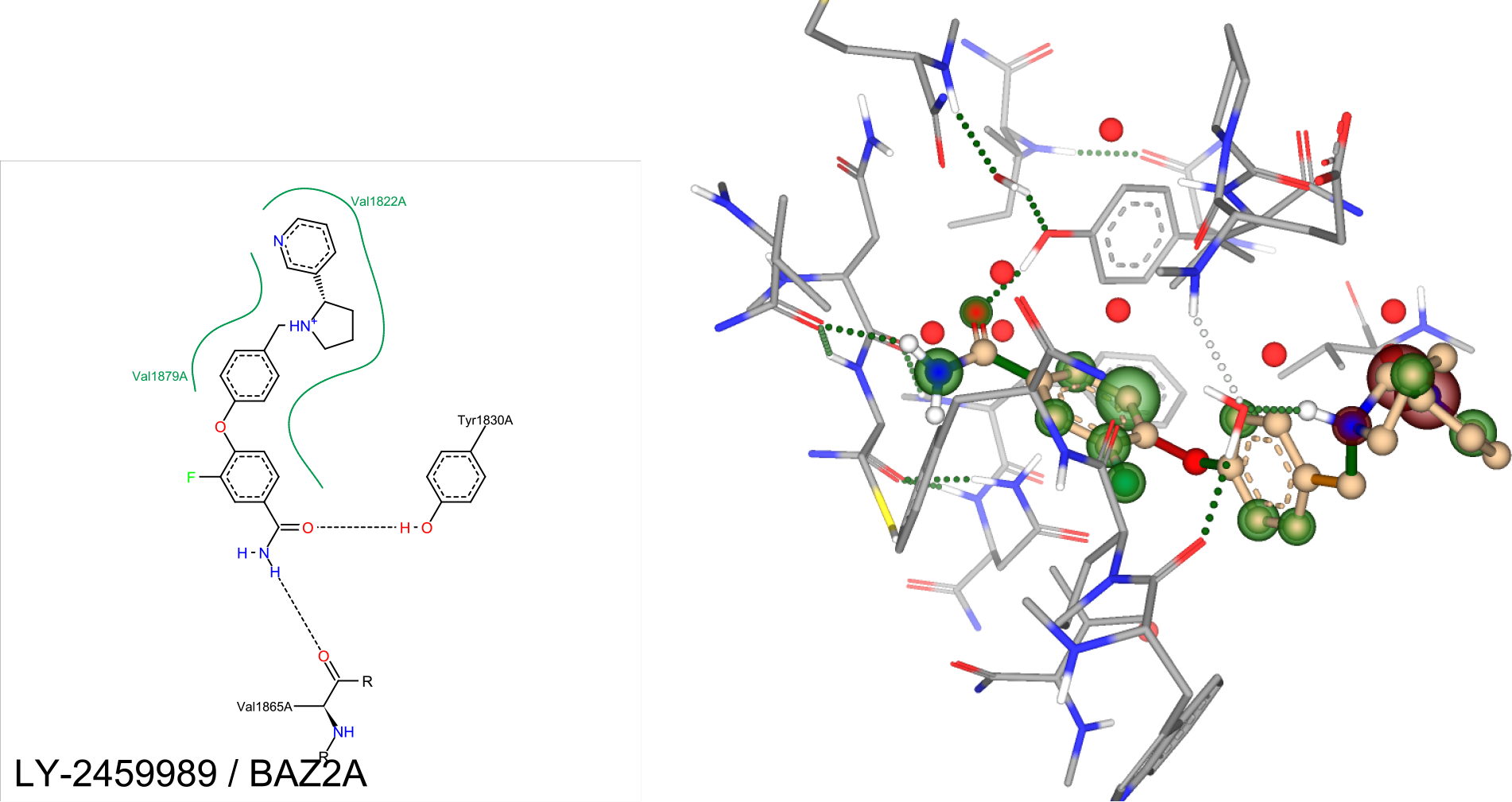
Three-dimensional docking view of LY-255582 against BAZ2A in SeeSAR

**Figure 19:**
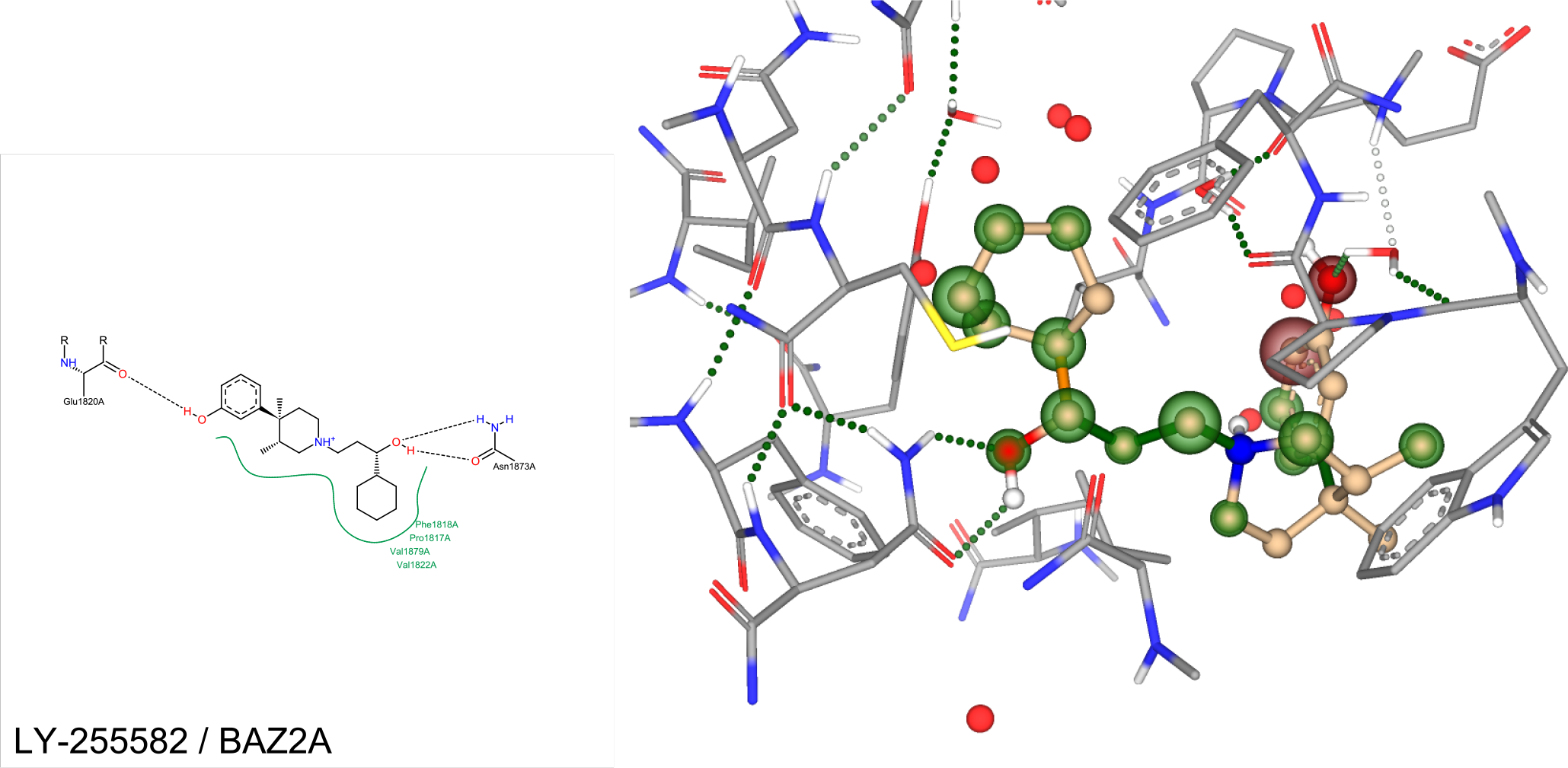
Three-dimensional docking view of LY-2459989 against BAZ2A in SeeSAR

#### Docking of LY-2459989 and LY-255582 against BAZ2B

Besides the estimated binding affinities being at least 40x weaker in comparison to PHIP(2), desolvation is the major contributor for in both cases, shown using distances between the ligands and the water molecules. In both cases, we get very disappointing results including many red coronas. This selectivity towards PHIP(2) over BAZ2B is good because PHIP(2) is category III and BAZ2B is category V.[14]

**Table 11:** Docking of LY-2459989 and LY-255582 against BAZ2B

| Ligand | HYDE<br>estimated<br>affinity lower<br>bound | HYDE<br>estimated<br>affinity upper<br>bound | H-bonds |
| --- | --- | --- | --- |
| LY-2459989 | 4.3 μM | 427 μM | 3 |
| LY-255582 | 73 μM | 7.3 mM | 2 |

**Figure 20:**
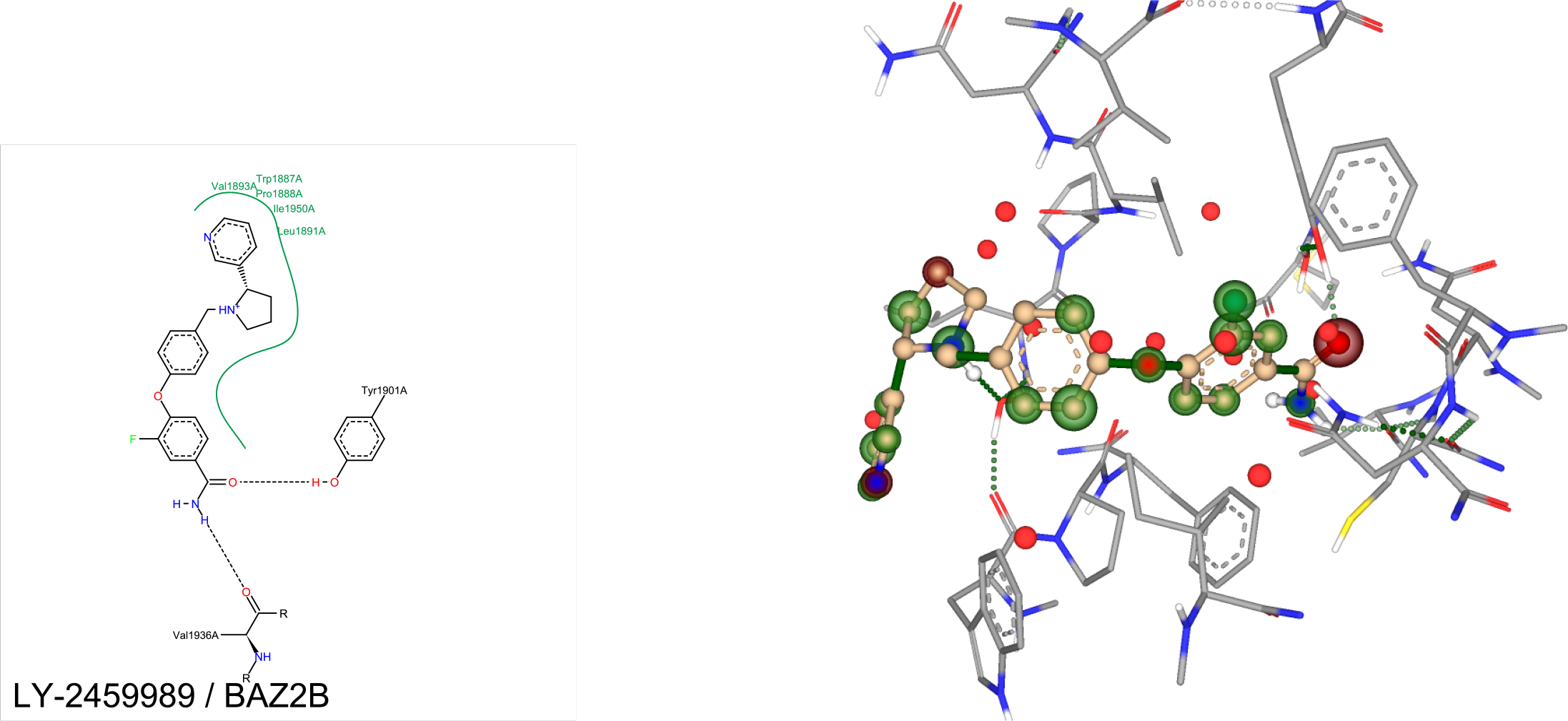
Three-dimensional docking view of LY-255582 against BAZ2B in SeeSAR

**Figure 21:**
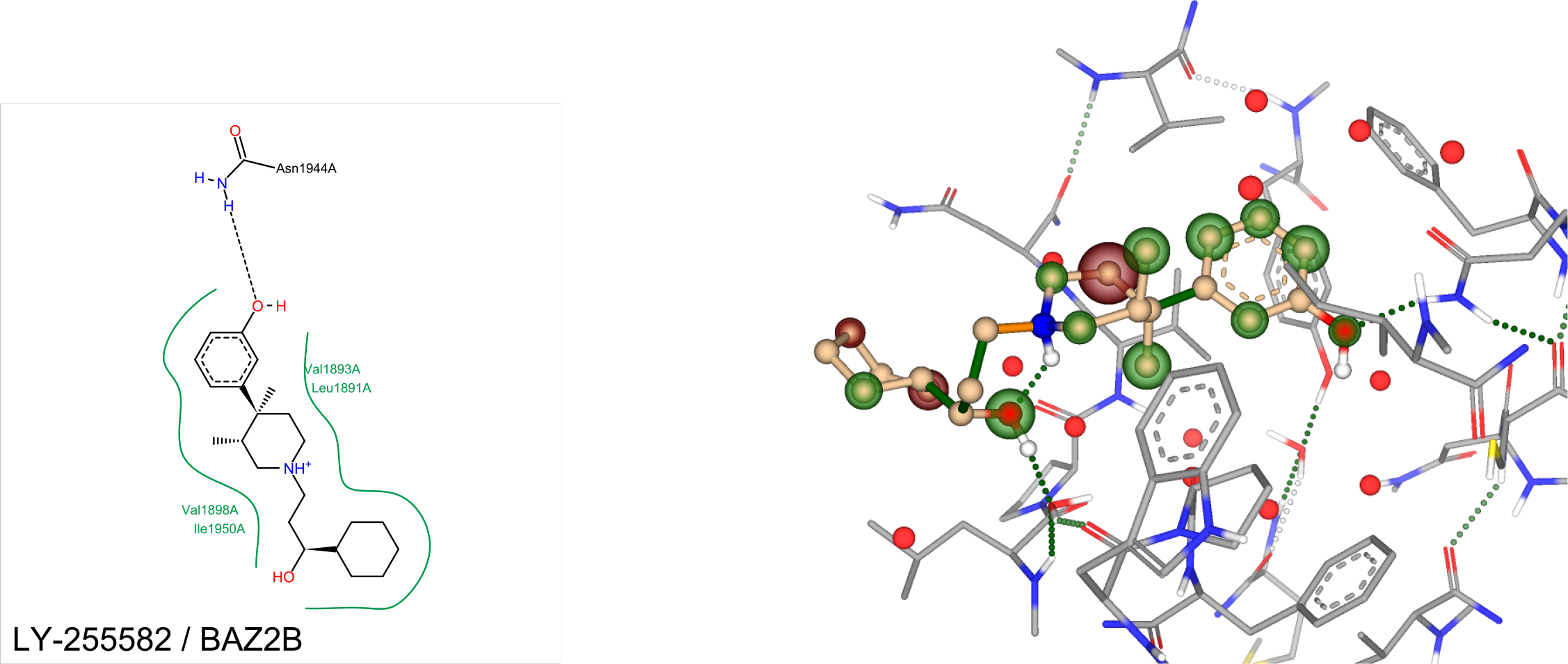
Three-dimensional docking view of LY-2459989 against BAZ2B in SeeSAR

#### Docking of published PHIP(2) hits against KOR PDB 4DJH

In order for there to be a link between KOR and PHIP(2), confirmed ligands for PHIP(2) must demonstrate good estimated binding affinity towards KOR, preferrably using the same residues being used to hold KOR antagonists in place. In this case, we chose PDB 4DJH chain A, which holds JDTic in place by interaction with residues VAL134, ASP138, MET142, and ILE294. Thus, we proceeded to dock all 7 currently published ligands, displaying the ligand of reference in chain A of PDB 4DJH[38], since that’s where binding with JDTic elicits an antagonistic response.

**Table 12:** Docking results of PHIP(2) ligands against KOR PDB 4DJH

| Ligand | HYDE<br>estimated<br>affinity<br>lower<br>bound<br>(KOR) | HYDE<br>estimated<br>affinity<br>upper<br>bound<br>(KOR) | $K_{iPHIP(2)}^{exp}$ | H-bonds | ASP<br>138 | MET<br>142 | ILE<br>294 |
| --- | --- | --- | --- | --- | --- | --- | --- |
| FMOOA463 | 142 nM | 14 μM | 68 μM | 1 | ✓ | ✓ | ✓ |
| XTS942 | 222 nM | 22 μM | 768 μM | 2 | ✓ | ✓ | |
| FMOOA322a | 1.6 μM | 161 μM | 190 μM | 2 | ✓ | | ✓ |
| FMOMB76b | 13 μM | 1.3 mM | > 5 mM | 1 | ✓ | | ✓ |
| FMOSA1515 | 28 μM | 2.8 mM | 269 μM | 0 | | | |
| FMOSA1544 | 90 μM | 9 mM | 588 μM | 2 | ✓ | ✓ | ✓ |

**Figure 22:**
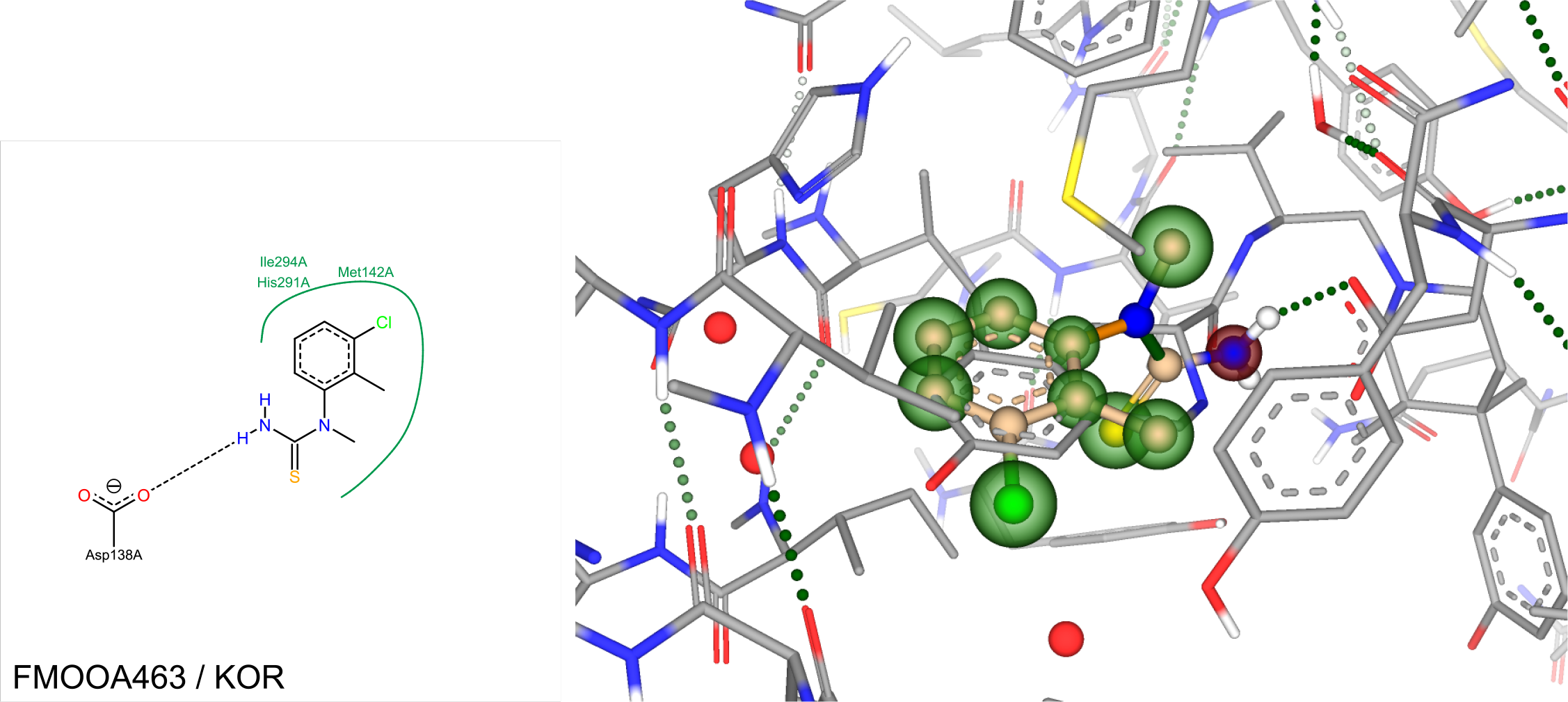
Three-dimensional docking view of FMOOA463 against KOR over JDTic as reference in SeeSAR

**Figure 23:**
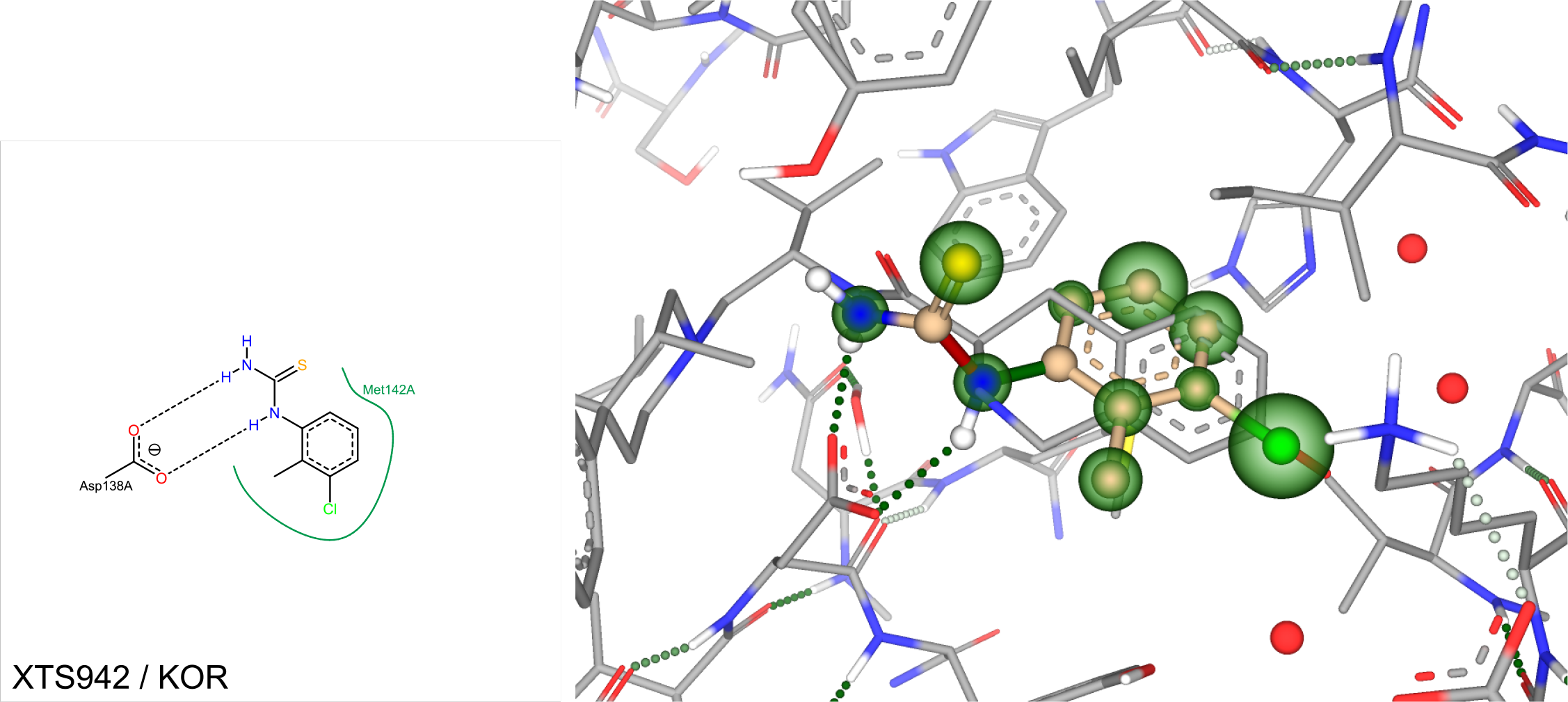
Three-dimensional docking view of XTS942 against KOR over JDTic as reference in SeeSAR

**Figure 24:**
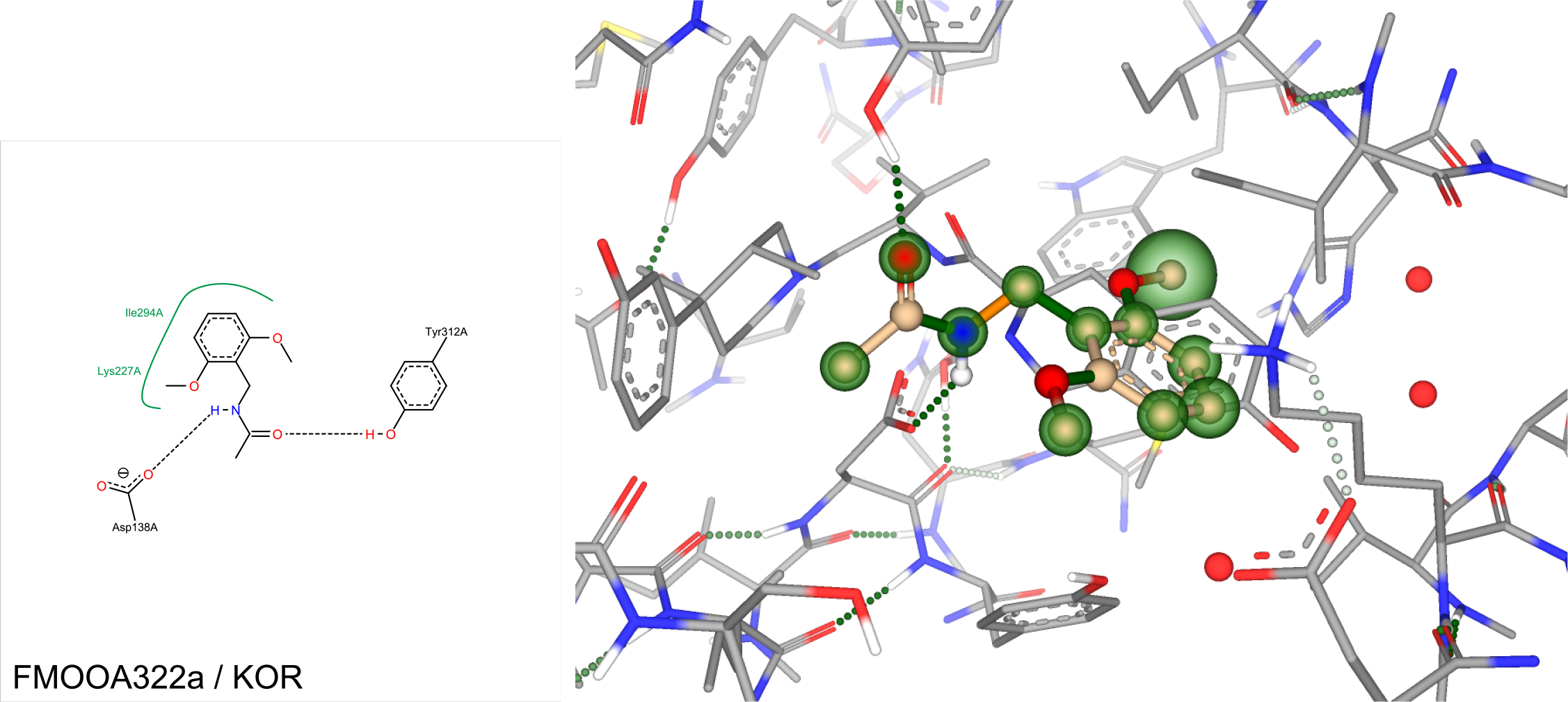
Three-dimensional docking view of FMOOA322a against KOR over JDTic as reference in SeeSAR

**Figure 25:**
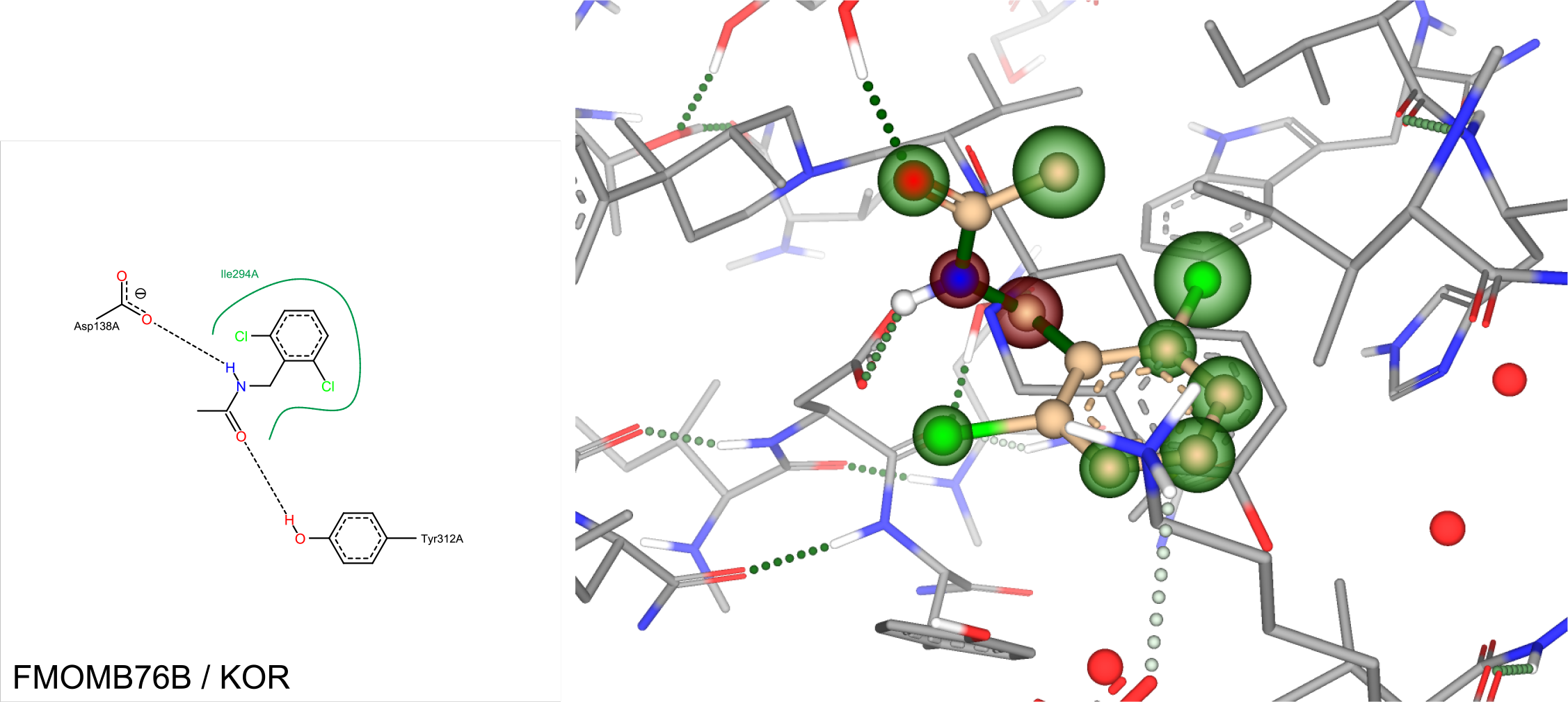
Three-dimensional docking view of FMOMB76b against KOR over JDTic as reference in SeeSAR

**Figure 26:**
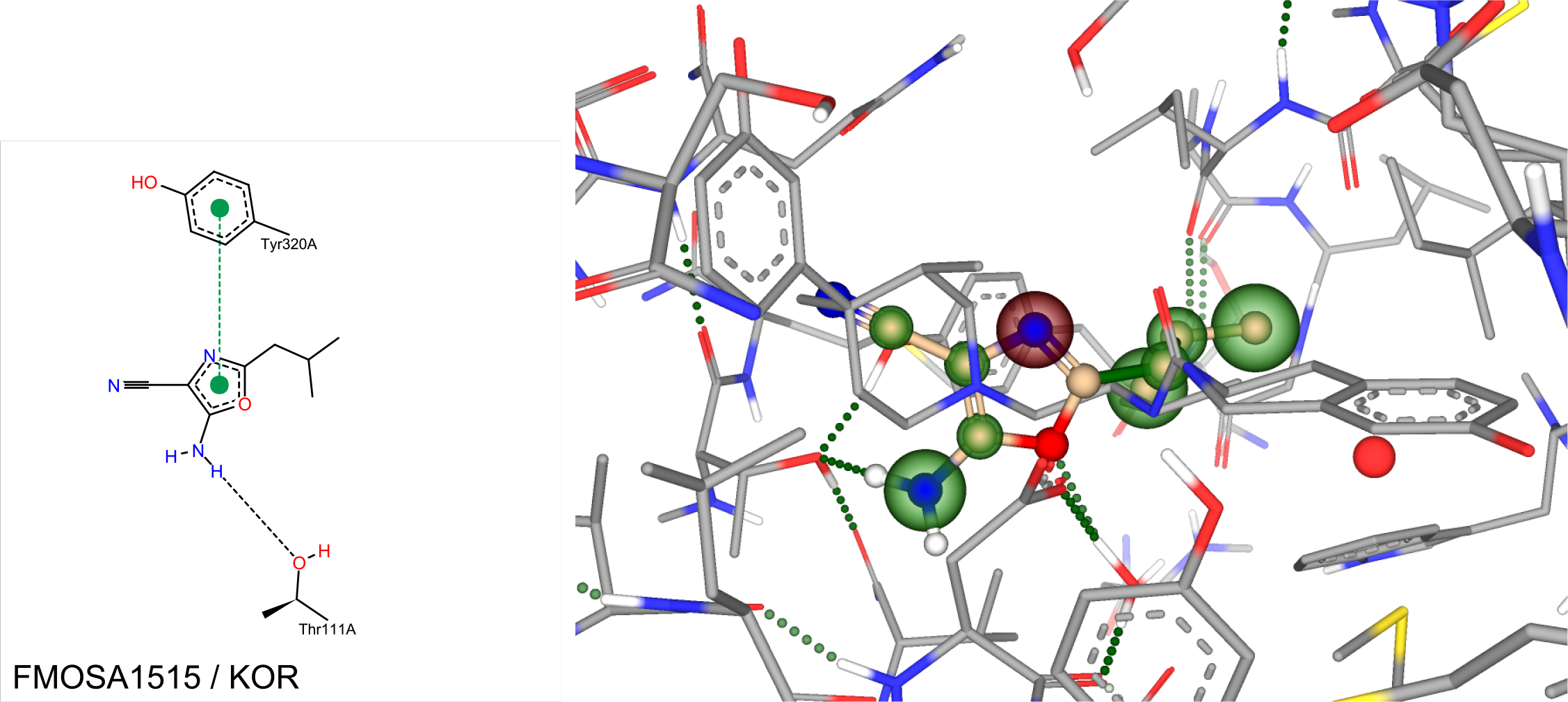
Three-dimensional docking view of FMOSA1515 against KOR over JDTic as reference in SeeSAR

**Figure 27:**
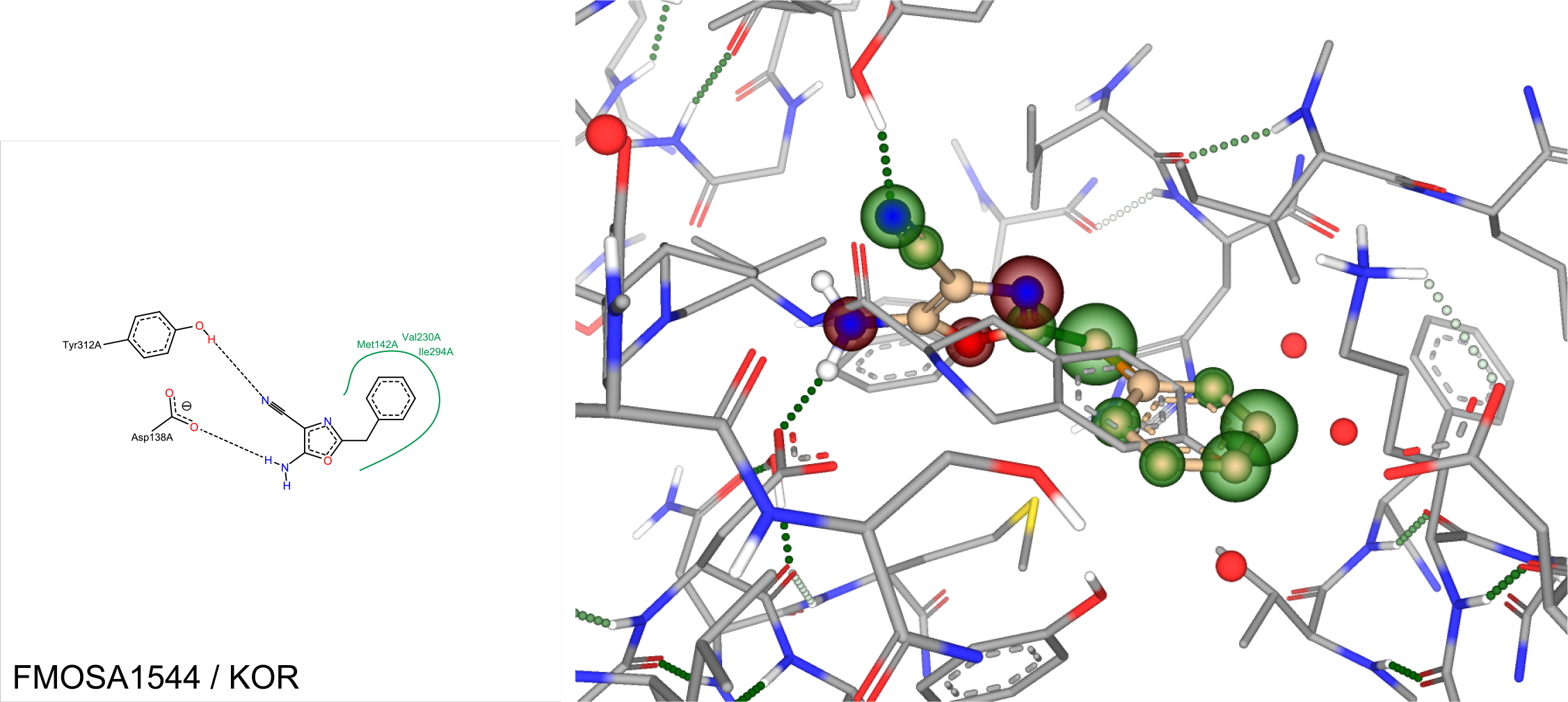
Three-dimensional docking view of FMOSA1544 against KOR over JDTic as reference in SeeSAR

These results are indeed very interesting, overlapping the original antagonist JDTic - in every single case, the estimated average binding affinity’s lower bound is much lower than the experimental values for PHIP(2). One should however note, that FMOSA1515 is predicted to not have any H-bonds towards KOR, and that an average between lower- and upper bound results in better binding towards PHIP(2) than KOR. Looking the average estimated binding affinity for each ligand, they are still much greater towards KOR in comparison to PHIP(2), except in the case of FMOSA1515 and FMOSA1544. The result for FMOSA1515 is however unreliable, since no H-bonds are predicted. ASP138A is the most occurring binding residue, followed by ILE294. In the case of XTS942, both nitrogens are predicted to form H-bonds towards ASP138A.

### Conclusion and future plans

Having found at least 2 new distinct pharmacophores with high estimated binding affinity towards PHIP(2) and with the exception of EP300, are weak binders at other bromodomains, which are strong binders at KOR/MOR, there could be a structure-activity relationship between two targets, in a similar way to how PHIP interacts with IRS1. In the future, we would like to test protein-protein interaction between bromodomains and Opioid receptors, as well as conduct binding studies and X-ray crystallography on the found ligands in combination with PHIP(2). Moreover, a relationship between PHIP(2)/EP300 and KOR antagonists could also mean that antagonists for other receptors are potentially targetting other bromodomains, opening the door for much more selective GPCR antagonist drug design, reducing unwanted side-effects as well as risk of great investment loss by late stage clinical trials failures.

### Methods

SeeSAR 8.0 and PoseView were downloaded from the BioSolveIT GmbH website (https://www.biosolveit.de/download/). The project was initialised by downloading the 5ENB PDB using the SeeSAR interface and mol files were created by copying the SMILES strings from the bindingDB website (https://www.bindingdb.org) and pasting them into MarvinSketch. These were imported into SeeSAR and every molecule was docked as follows:

1. Generate poses
2. Calculate estimated affinity
3. Removal off all but the highest affinity pose
4. Repeat steps 1 through 3 until the suffix of the highest binding ligand is shorter than that of all of the other poses for the same ligand

Once this was done for all molecules, each ligand was exported individually and PoseView was used with the following command: poseview.exe -l <ligand>.sdf -p 5enb.pdb -o <ligand>.svg -t <ligand>

## Acknowledgements

The author thanks BioSolveIT GmbH (especially Dr Marcus Gastreich), Dr Evet Ghobrial, Dr Diana Jordan, and Dr Qiao Chen for their support throughout the years.

